# A cooperative switch within the KaiC hexamer revealed by cryo-EM

**DOI:** 10.1101/2022.02.27.481910

**Authors:** Xu Han, Dongliang Zhang, Lu Hong, Daqi Yu, Zhaolong Wu, Tian Yang, Michael Rust, Yuhai Tu, Qi Ouyang

## Abstract

The circadian clock of cyanobacteria is based on an approximately 24h rhythm in the phosphorylation level of KaiC, a hexameric ATPase. This oscillation can be reconstituted *in vitro* by incubating three proteins together with ATP. Like all chemical oscillators, this system must include a nonlinear, or switch-like, feedback loop, whose nature has been unclear. Here, by using single particle cryo-EM at near-atomic resolution we identified two major conformational states of KaiC subunits, denoted as the exposed state and the buried state, which may provide a structural basis of how the KaiC hexamer changes its conformation during the (day-night) phosphorylation-dephosphorylation cycle. We classify the abundance and pattern of exposed and buried states within hexamers for more than 160,000 KaiC particles. The statistics of the spatial arrangement of the two states in hexamers can be quantitatively fit by a simple statistical physics model with an interaction energy between neighboring subunits and a local field that depends on phosphorylation state. Our study shows that phosphorylation shifts the probability of each conformation and reveals that there is substantial cooperativity between neighboring subunits, which can allow a KaiC hexamer to respond in an ultrasensitive, switch-like manner to changes in the phosphorylation level.

## Introduction

Circadian clocks are endogenous biological processes that exhibit self-sustained oscillations with an approximately 24h period^1,2^. These rhythms are found across diverse organisms from prokaryotes to eukaryotes^3,4^. In animals, disruption of circadian clock function causes temporal disorganization of physiology and contributes to a variety of diseases, such as disorders of the nervous system, cancer, and cardiovascular and cerebrovascular diseases^5–9^.

Cyanobacteria are the simplest organisms known to possess a circadian clock^10,11^. Because the cyanobacterial oscillator can be reconstituted *in vitro* using three proteins, KaiA, KaiB and KaiC^12^, it presents an opportunity to uncover fundamental biophysical principles of circadian timing. This post-translational oscillator is realized by the interactions and conformational changes among the three Kai proteins, and the oscillation is manifest as rhythmic phosphorylation of KaiC ^12,13^.

The key enzyme in this circadian clock is KaiC. It forms a homo-hexamer with a “double doughnut” shape where each subunit consists of two tandem RecA-superfamily ATPase domains, named the N-terminal CI domain and C-terminal CII domain, with a total of 12 nucleotides bound to each hexameric particle^14,15^. Both the CI domain and CII domain have ATPase activity, and CII domain also possesses a phosphotransferase activity allowing the protein to autophosphorylate and dephosphorylate^16–21^. KaiC has two observed phosphorylation sites in the CII domain, Ser431 and Thr432, whose phosphorylation and dephosphorylation follow a cyclic order: ST→ SpT→ pSpT→ pST→ ST^22,23^. During the subjective day, KaiA stimulates phosphorylation by binding to the KaiC C-terminal tail and remodeling the A-loop, allowing nucleotide exchange ^19,24–29^.

This process is terminated in the night phase of the cycle when KaiB binds to phosphorylated KaiC and sequesters KaiA, allowing KaiC to dephosphorylate^30–35^. While this describes changes that occur to an individual KaiC hexamer, for the entire reaction to oscillate coherently, mathematical modeling shows that the transition between high KaiA activity and low KaiA activity must be switch-like or ultrasensitive. The molecular mechanisms of this proposed ultrasensitivity are still unclear. Because the phosphorylation of KaiC both stores information about the time of day and determines the level of KaiA inhibition, a major question is how phosphorylation alters the structure of KaiC. This question has been difficult to address. Previously reported high-resolution crystal structures with different phosphorylated and phosphomimetic states are nearly identical ^14,15,26,35,36^, suggesting that there are functionally important conformational states of KaiC present in solution that are difficult to capture using crystallography.

Indeed, previous studies suggest that KaiC has dynamic structural properties that change across a phosphorylation cycle^29,36–38^. The flexibility of KaiC CII ring and the status of the A-loops are determinants of KaiC phosphorylation activity^28,36^. In principle, cryo-electron microscopy (cryo-EM) can not only resolve static protein structures, but also could be a powerful tool to study the dynamic interconversions of proteins in solution. Here, we report our investigations of statistical characteristics of the conformations of KaiC. Two KaiC mutants were used in this study. One mimics the dawn-like dephosphorylated state via alanine substitution at the phosphorylation sites (KaiC-AA), the other mimics the dusk-like fully phosphorylated state via glutamate substitution (KaiC-EE). These mutants have been previously shown to differentially allow KaiB binding^22,39^, indicating that they simulate important features of KaiC at different times of day.

By classifying individual KaiC subunits, we found that the status of the A-loop could be treated approximately as a two-state system—some subunits have an A-loop that is largely buried in the subunit interface (buried state), and others have an extended, flexible A-loop protruding from the KaiC particle (exposed state). We find that these two states exist in a dynamic equilibrium because both states can be found in each mutant, suggesting that KaiA stimulates KaiC through a conformational selection mechanism. The buried state is much favored in the KaiC-EE mutant and we occasionally observed a distinct structure where the buried A-loop makes direct contacts with the 422-loop, previously implicated in the regulation of the phosphorylation reaction^40^.

We used an Ising model to describe correlations of the conformational states found within each hexamer^41–43^. This analysis reveals that there is substantial nearest-neighbor cooperativity in the ring with adjacent subunits tending to adopt the same conformation. This model gives quantitative agreement with the observed frequency of hexamers with different patterns of conformational states. The picture that emerges is one where the A-loops within a KaiC hexamer form an intrinsically cooperative switch, and the role of phosphorylation is to bias the switch towards the exposed or buried state.

## Results

### Cryo-EM structure determination of KaiC-AA and KaiC-EE

To investigate the effects of different phosphorylation states on structures and functional mechanisms in Kai system, two KaiC phosphomimetic mutants were used (KaiC-AA (S431A, T432A) and KaiC-EE (S431E, T432E)). These two mutations mimic the state of the clock near dawn and near dusk, respectively. We used a FEI Titan Krios G2 microscope device to collect cryo-EM data of KaiC-AA and KaiC-EE after incubation in the presence of 1mM ATP at a functionally relevant state (see Methods).

We first focused on two structures: one in KaiC-AA (Fig. 1A) and one in KaiC-EE (Fig. 1B), both refined to nominal resolution of 3.3 Å with relatively clear secondary structures (Fig. 1C, 1D, Extended Data Fig. 1). These two cryo-EM densities (Fig. 1A, 1B) were superimposed together for comparison (Extended Data Fig. 2A). We found that the KaiC-EE density is more compact in the CII domain. This observation is consistent with previous reported Trp fluorescence results showing that the overall shape of KaiC is more loosely packed in S/T state than in pS/pT state^37^. The difference map (Extended Data Fig. 2B) calculated by RELION^44,45^ indicates that this density corresponds to a pair interacting loops, *i.e*., the A-loop (residue 488-497) and the 422-loop (residue 417-429). In the refined KaiC-EE atomic model, distance between G421 (belongs to the 422-loop) and S491 (belongs to the A-loop) is about 3.8 Å (Extended Data Fig. 2C).

**Fig. 1.**
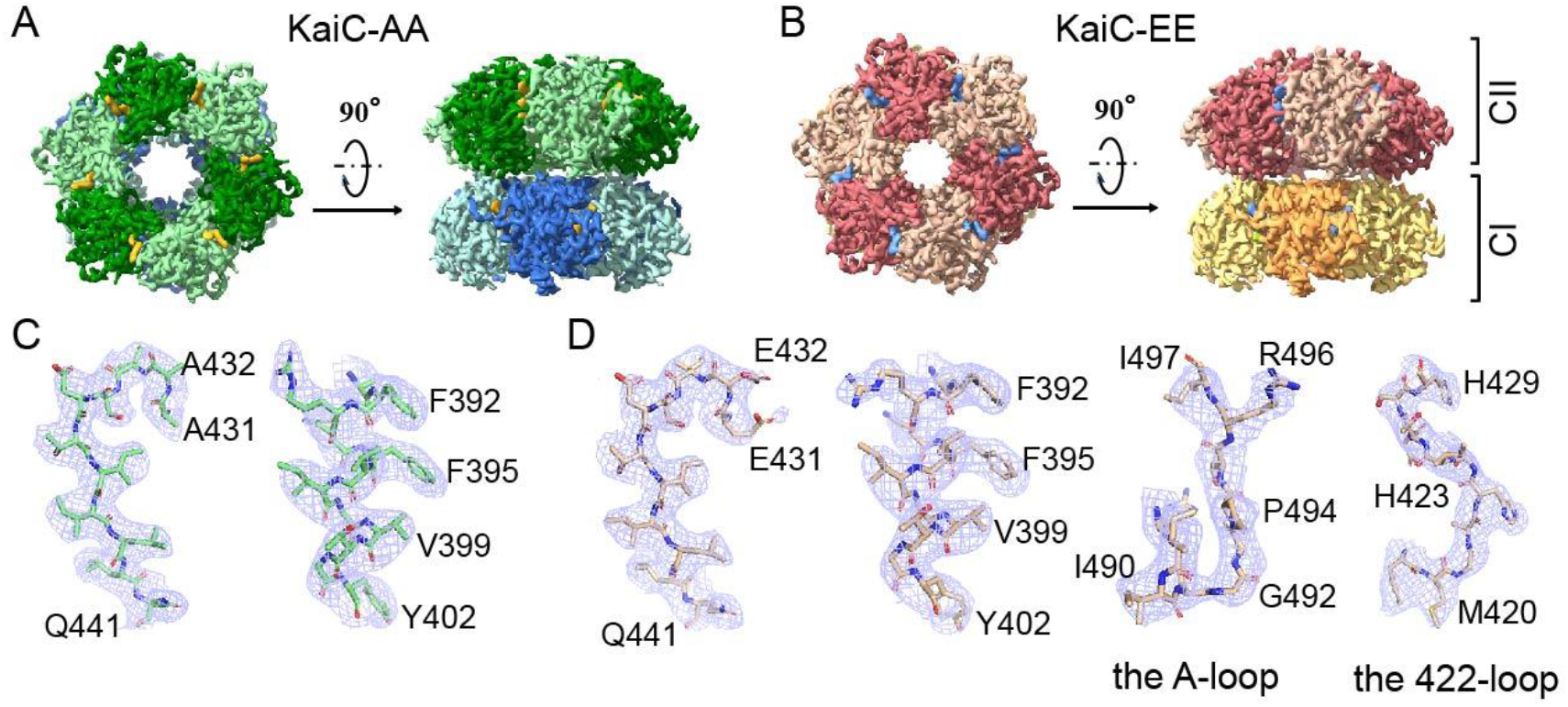
Cryo-EM maps of KaiC-AA and KaiC-EE. (**A**) Top and side view of KaiC-AA, with CII and CI domains are colored in dark green and green and in blue and light blue, respectively. (**B**) Top and side view of KaiC-EE, with CII and CI domains are colored in red and pink and in orange and yellow, respectively. (C) and (D) show typical high-resolution densities of the secondary structures in the cryo-EM structure of KaiC-AA (**C**) and of KaiC-EE (**D**).

### Two functional conformations for each KaiC subunit in a hexamer

To further study the dynamic behaviors of KaiC-AA and KaiC-EE, we focused on CII domain using six-fold pseudo-symmetry (see Methods) by RELION^44,45^. There are two distinct possible conformations (exposed and buried) for each KaiC subunit in a KaiC hexamer (Fig. 2A). For the exposed state, the A-loop tends to stick out of the hexamer and is very dynamic, thus most of them do not have clear density. For the buried state, the A-loop forms a well-defined “U-shaped” line inside the central channel; this stable conformation contributes to a strong density in the electron density map. Previous work has been proposed that hydrophobic, electrostatic, and hydrogen bond interactions exist in the regions where KaiA and KaiC interact^24^. The A-loop in the exposed state is likely more available to contact with KaiA.

**Fig. 2.**
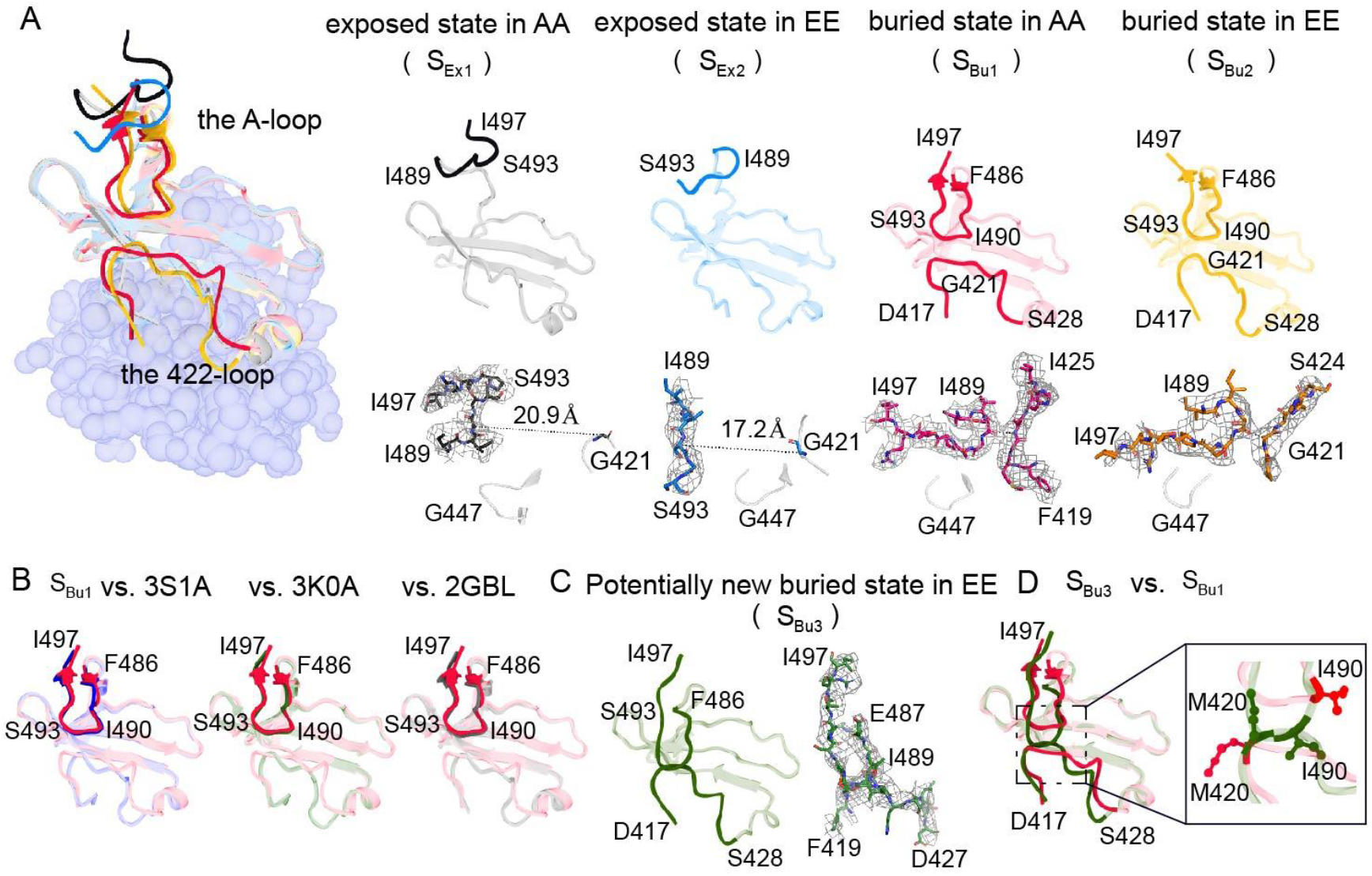
Different conformational states of the A-loop and 422-loop area. (**A, Left**) The overview diagram of typical exposed and buried state in KaiC-AA and KaiC-EE. (**Right**) Corresponding independent exposed and buried state in KaiC-AA and KaiC-EE (denoted as *S*_*Ex*1_, *S*_*Ex*2_, *S*_*Bu*1_, *S*_*Bu*2_, colored in black, royal blue, red, orange, respectively). At the bottom are the close-up view of the A-loop shown in a stick representation superimposed over its corresponding cryo-EM densities in a gray mesh at 3σ, 5σ, 9σ, 9σ level representation, respectively. (**B**) The buried state (*S*_*Bu*1_) superimposed with previously reported crystal structures in PDB data bank (with 3S1A, 3K0A, 2GBL colored in blue, green and dim gray, respectively). The A-loop is shown as cartoons without transparency. (**C**) A potentially new buried state conformation only found in KaiC-EE data set (denoted as *S*_*Bu*3_, colored in dark green), with the A-loop and 422-loop densities shown as a gray mesh at 11σ level on the right. (**D**) Superimposed the atomic model of *S*_*Bu*1_ and *S*_*Bu*3_, with side chains of M420 and I490 are given in a close-up view and labeled.

We also compared the structure of the buried state with previously reported crystal structures ^14,15,26,46,47^ and found that all structures were almost identical in the A-loop area (Fig. 2B, Extended Data Fig. 3). Thus, it seems reasonable that the crystal packing forces these crystal structures into what we call the buried state. However, it is worth noting that there is another type of buried state only observed in KaiC-EE data set, *i.e*., a potentially new buried state in which the A-loop and 422-loop are directly connected (Fig. 2C). This is mainly due to movements of two residues: the Met420 side chain moves up while the Ile490 side chain moves down (Fig. 2D). This potentially new conformation indicates a stronger interaction between the A-loop and 422-loop, which is consistent with the idea that the A-loop restrains the motion of 422-loop as previously reported^40^.

### Strong conformational cooperativity in KaiC hexamers

By using RELION^44,45^ to classify the original cryo-EM particles (1,592,573 KaiC-AA particles and 934,373 KaiC-EE particles), we selected a subset of particles with high quality well-defined structures (140,475 particles for KaiC-AA and 371,557 particles for KaiC-EE). All the KaiC monomers in these selected hexamer particles are then refined and clustered into a number of clusters (16 used here), each of which is represented by the averaged structure (3D volume) of the individual particles belonging to the cluster. (See Extended Data Figs. 4 and 6 for the data processing flowcharts for KaiC-AA and KaiC-EE respectively.) Next, each KaiC monomer cluster is classified as the buried (Bu) or exposed (Ex) conformational state based on the overlap between its 3D volume and two structure masks that characterize the Bu conformation (see Extended Data Figs. 5 and 7 for details.) As a result of this analysis, we can assign each monomer in all the selected hexamers one of two states: Bu or Ex. There is a small fraction of monomers (25.7% for KaiC-AA and 18.2% for KaiC-EE) that can not be classified as either Bu or Ex with sufficient statistical confidence. We call their conformation Undefined (Un). These Un monomers do not have a well-defined structure. They may represent the transitional state(s) between the Bu state and the Ex state or they could be caused by inaccuracy in our experiments.

We first studied the statistics of the conformational states of the KaiC monomers. We found that the probabilities of KaiC monomers being in the exposed or buried state depends on its phosphorylation state. KaiC-AA is more likely to be in the exposed state, whereas KaiC-EE is more likely to be in the buried state (Extended Data Figs. 4 and 6). Next, we investigated the statistics of the conformational states of the 6 subunits (monomers) in a hexamer. For a KaiC hexamer, there should be 2^6^ = 64 possible configurations (arrangements) of the 6 monomers, each with two possible conformational states. Considering the rotational degeneracy, these configurations can be combined into 13 conformational patterns each with a degeneracy index *Ω_k_ (k=1, 2,…, 13)*, which corresponds to the number of configurations that pattern-*k* contains: 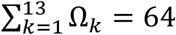 (see Supplementary Note 1 and Supplementary Fig. 1 for details). We put the conformational state of the monomers back into their positions in hexamers and counted the probabilities of these 13 conformational patterns for KaiC-AA (Fig. 3A) and KaiC-EE (Fig. 3B), respectively, for those hexamers in which all 6 monomers have clearly defined conformational states (24,240 particles for KaiC-AA, 116,785 particles for KaiC-EE). To avoid bias in particle selection, we also performed a statistical analysis including a large number of particles not used in the analysis shown in Fig. 3 due to orientation preference. These statistical results indicated that including these particles did not change the results (see Supplementary Note 1 and Supplementary Fig. 2).

**Fig. 3.**
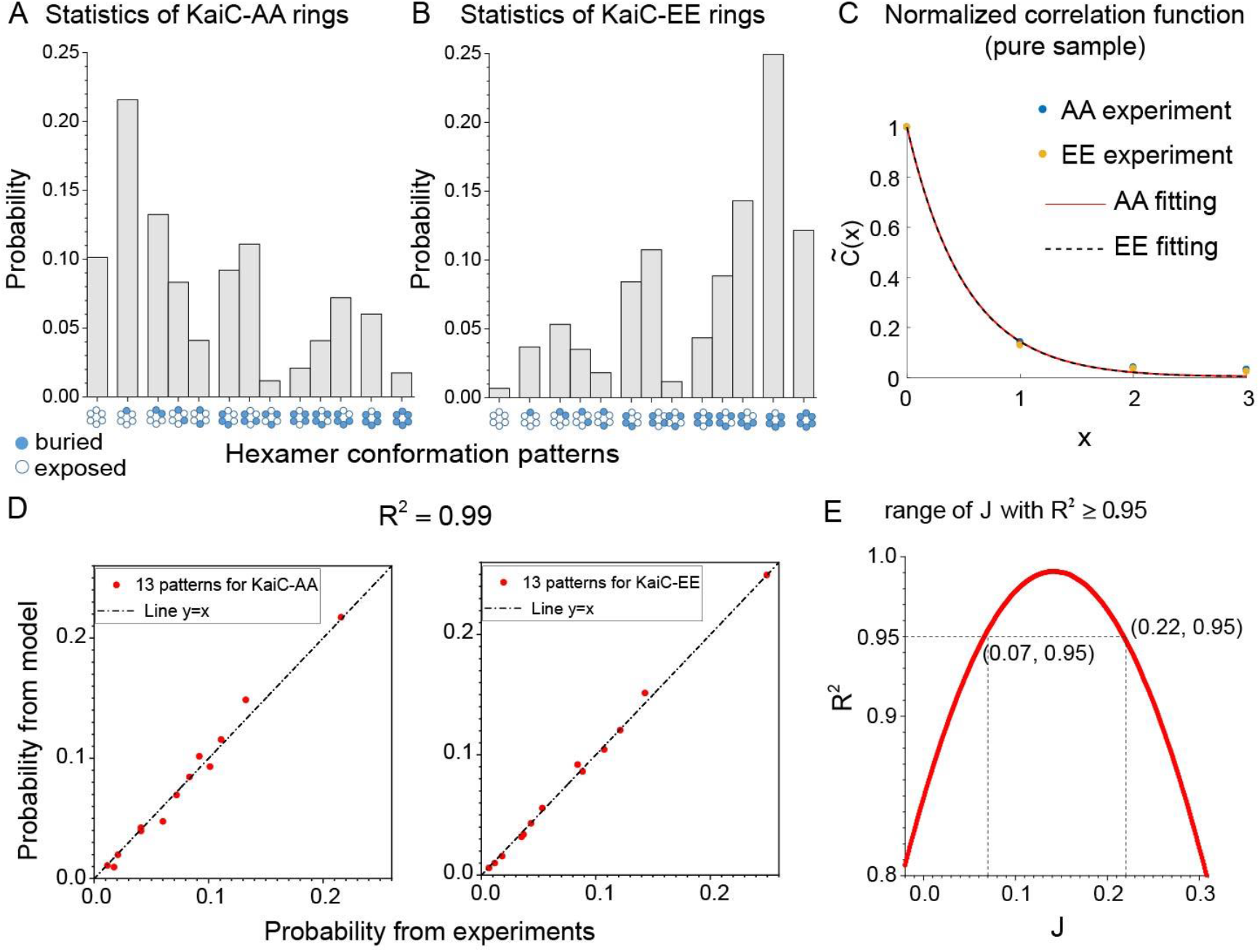
Statistical results for pure KaiC-AA and KaiC-EE hexamers. Probabilities of KaiC hexamers in different conformational patterns for KaiC-AA data set (**A**) (24,240 rings) and for KaiC-EE data set (**B**) (116,785 rings). The patterns are grouped and ordered according to the total number of exposed (or buried) states in the hexamer (total “spin”). Given the large number of hexamers in each pattern (>100 particles), the relative errors are small (< 10%). (**C**) Normalized correlation function for KaiC-AA and KaiC-EE hexamers, respectively. Dots are calculated from experimental data whereas lines are the theoretical results from the Ising model. About 10^6^ subunit pairs are used to compute 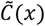, so the error is ~10^-3^, which is too small to show in the figure. **(D)** Comparison between experiment and model for KaiC-AA (left) and KaiC-EE (right), respectively. **(E)** Optimal *R*^2^ varies with coupling constant *J*. In the range of *J* ∈ [0.07, 0.22], *R*^2^ is higher than 0.95, with the maximum at *J* = 0.14.

Qualitatively, this analysis (Fig. 3A, 3B) reveals that there is cooperativity in the conformational transitions of KaiC hexamer. Indeed, if all monomers in a hexamer were independent of each other, the probability distribution of the hexamer conformational patterns would follow a simple binary distribution: *P_k_* = *Ω_k_p^nk^*(1 – *p*)^6–*n_k_*^, where *p* is the probability of a subunit in the exposed conformational state and *n_k_* is the number of exposed state in hexamer pattern-*k* (see Supplementary Fig. 1B for values of *Ω_k_* and *n_k_* for *k* = 1,2,…, 13). It is easy to see from our data that this is not the case. For example, the two hexamer conformational patterns ‘Ex-Ex-Ex-Ex-Bu-Bu’ (k=3) and ‘Ex-Ex-Ex-Bu-Ex-Bu’ (k=4) have the same Ω_3_ = Ω_4_ = 6 and *n_3_* = π_4_ = 4, which would lead to their probabilities being equal without KaiC-KaiC cooperativity. However, our experiment results showed that *P_3_ > P_4_* for both KaiC-AA and KaiC-EE (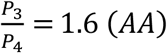, 1.5 (*EE*)), which clearly indicates positive cooperativity among individual monomers that favors the neighboring monomers to be in the same conformational state.

To quantify this cooperativity, we calculated the pair-wise (subunit-subunit) conformational correlation function for each dataset (EE or AA):

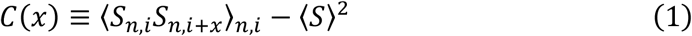

where *i* (= 1,2,…,6) is the ordered subunit index with periodic boundary condition (*S*_*i*+6_ = *S_i_*) and *n* represents different hexamers in the dataset. The state variable *S_n,i_* of subunit-i in the hexamer-n can be “+1” or “-1”, corresponding to an exposed state or a buried state, respectively. ⟨S⟩ is the average state variable over all monomers in either Bu or Ex states, and the average ⟨*S_n,i_S_n,i+x_*⟩_*n,i*_ is taken over all hexamers (*n*) and all pairs of monomers (*i*, *i* + *x*) in a hexamer except for those in which at least one of the monomers in the pair has an undefined conformation (Un). As is shown in Fig. 3C, the normalized correlation function 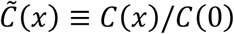 is significantly larger than zero for *x* ≥ 1, which clearly indicates subunit-subunit cooperativity.

### An Ising model quantitatively explains the experimental data

Next, we explain the observed statistics of the conformational states in KaiC hexamers (Fig. 3A-C) by using a simple model that includes cooperativity among monomers in a hexamer. Given the individual conformational state of each monomer in hexamers, the all-or-none Monod-Wyman-Changeux (MWC) model^48^ certainly does not fit the experimental data. We thus adopted the more general Ising-type model^49^, which is a minimal model that considers only the most salient features suggested by experiments: 1) the conformation of a subunit depends on its phosphorylation level; 2) there is a positive cooperativity between neighboring subunits in the hexamer.

In the minimal model, only the nearest neighbor subunits interact with each other, and the Hamiltonian (energy) of each configuration (64 in total) can be expressed as:

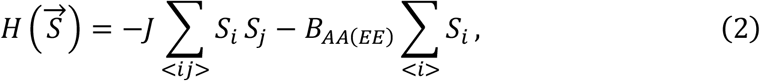

where *∑*⟨*ij*⟩ represents a sum over all nearest neighboring pairs. *J* is the coupling constant—a positive value of *J* favors the two neighboring subunits to be in the same conformational state; *B_AA_* (or *B_EE_*) is the “local field” for the KaiC-AA (or KaiC-EE) subunit—a positive (negative) local field favors the exposed (buried) state for the subunit. Assuming the system is at thermodynamic equilibrium, the probability of each hexamer conformational pattern can be expressed as:

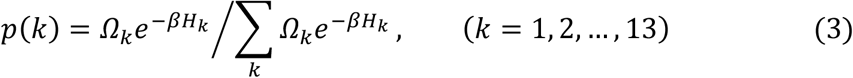

where *H_k_* is the energy for hexamer pattern-*k*, and 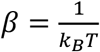 with *k_B_* the Boltzmann constant and *T* an effective temperature. The effective thermal energy is set to be the energy unit (*k_B_T* = 1) henceforth. Fig. 3D demonstrate the best fit of the experimental data to our model. *R^2^* between the model prediction and the experimental data is 0.99 (Fig. 3D, consider both KaiC-AA and KaiC-EE together) indicating that the experimental data can be quantitatively described by a simple Ising model with only nearest neighbor interactions. In Fig. 3E, we plot the dependence of *R^2^* on the coupling constant *J*, which clearly shows that our data cannot be explained without cooperativity (*R*^2^=0.84 when *J*=0). The best parameters for fitting both the KaiC-AA and KaiC-EE mutant data are: *J* = 0.14 ± 0.07, *B_AA_* = 0.19 ± 0.04, *B_EE_* = −0.25 ± 0.04 (error bars computed with *R^2^* ≥ 0.95, see Fig. 3E). These parameters indicate that there is substantial cooperativity between nearest neighbor subunits, and different local fields caused by Alanine or Glutamate mutation have opposite effects on the propensity of the exposed state or buried state.

The normalized correlation function can be determined exactly in the Ising model:

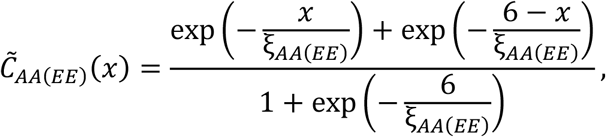

where the correlation length for KaiC-AA and KaiC-EE are respectively: 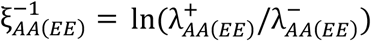, with 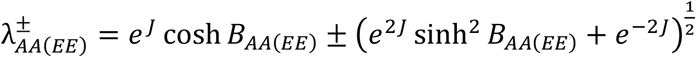. We found that the correlation function from the Ising model fits the observed correlation accurately (Fig. 3C). The best fit parameters, *J* = 0.15, *B_AA_* = 0.21, and *B_EE_* = −0.22, are in quantitatively agreement with those obtained from fitting the configuration data (Figs. 3A, 3B, 3D). The correlation length is found to be short: ξ*_AA_* ≈ ξ*_EE_* ≈ 0.5 < 1, which confirms that the dominant subunit-subunit interactions are those between neighboring subunits in the KaiC hexamer.

Notice that the coupling constant *J* is the same in KaiC-AA and KaiC-EE, which means that the Ala or Glu mutation only affects the propensity of each individual subunit, rather than subunit-subunit cooperativity. In a previous study^39^, the subunit free energy difference between two functional states (competent and incompetent to interact with KaiB) is estimated to be 1 ± 0.14 for unphosphorylated (ST) KaiC and −1 ± 0.69 for doubly phosphorylated (pSpT) KaiC based on modeling the observed oscillatory dynamics of the KaiC system. From the Ising model studied here, the free energy difference between the exposed state and the buried state for a single subunit is Δ*H_AA_* = 2*B_AA_* = 0.38 ± 0.08 and Δ*H_EE_* = 2*B_EE_* = −0.5 ± 0.08 for the AA (mimicking ST) and EE (mimicking pSpT) mutants, which are consistent with the previous estimates. The quantitative difference may be due to differences between mutant (AA and EE) and wild-type (ST and pSpT) proteins.

### Cooperativity in hexamers with heterogeneous phosphorylation

In an oscillating reaction, KaiC hexamers likely contain mixtures of differently phosphorylation subunits. To test the generality of cooperative interactions between neighboring KaiC monomers in the hexamer, we constructed a mixed sample with both KaiC-AA and KaiC-EE monomers^21^ and measured the distribution of KaiC conformational states in hexamers. Briefly, KaiC mutants (KaiC-AA, KaiC-EE, 1:1) were buffer exchanged into the running buffer with 0.5 mM ADP, and incubated at 4°C for about 24 hours to disrupt hexamer structure. Then monomerized KaiC-AA and KaiC-EE were mixed before re-hexamerization via the addition of ATP^39^. We collected cryo-EM data of this mixed sample with FEI Titan Krios G2 microscope (Extended Data Fig. 8A). After unsupervised 2D classification (Extended Data Fig. 8B) and 3D refinement by RELION^44,45^, we obtained the cryo-EM density map that was refined to nominal resolution of 3.8 Å (Extended Data Figs. 8C and D). By following the same procedure for the pure samples, we obtained the probabilities of the 13 hexamer conformational patterns for the mixed sample as shown in Fig. 4A.

**Fig. 4.**
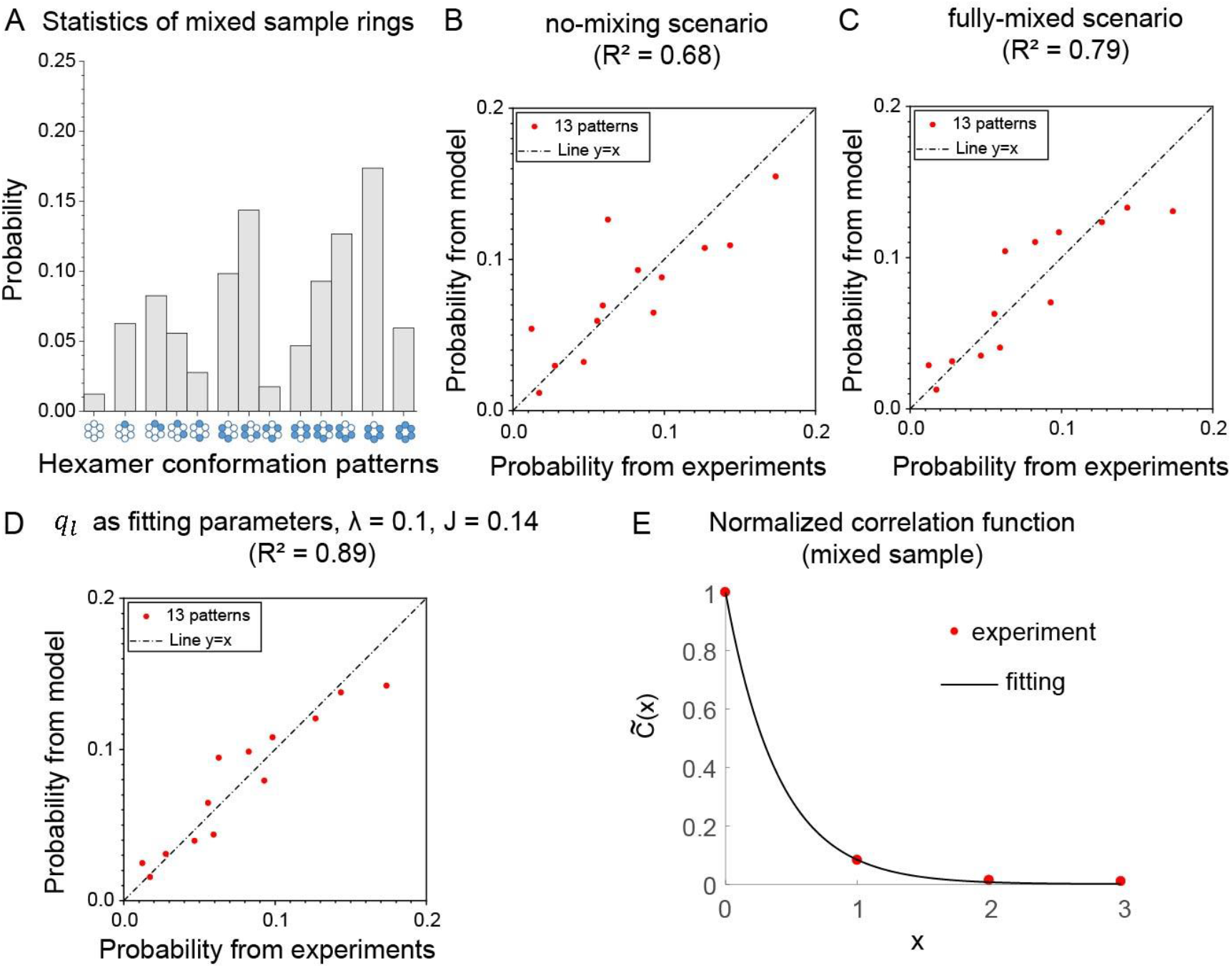
Statistical results for the mixed hexamers. (**A**) Probabilities of KaiC hexamers in different conformational patterns for the mixed sample data set (19,858 rings). Given the large number of hexamers in each pattern (>100 particles), the relative errors are small (< 10%). Comparison between the mixed experimental data and model results with the no-mixing scenario (**B**), the fully-mixed scenario (**C**), *q_l_* treated as free parameters and *λ* = 0.1, *J* = 0.14 (**D**). (**E**) Normalized correlation function of the mixed sample. Dots are calculated from the experimental data while the solid line is from the fitting by the Ising model. About 10^6^ subunit pairs are used to calculate 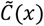, so the error is ~10^-3^, too small to show in the figure. The fitting parameters are: *J_mix_* = 0.086, *B_mix_* = −0.16.

In the mixed sample, each hexamer can have 14 distinct KaiC-AA and KaiC-EE subunit arrangements (see Supplementary Note 2 and Supplementary Table 1); the probability of these subunit arrangements is denoted as *q_l_*(*l* = 1, 2,…,14). The Hamiltonian of different hexamer conformational pattern *k*(*k* = 1, 2,…,13) can be determined by the Ising model for arrangement-*l*:

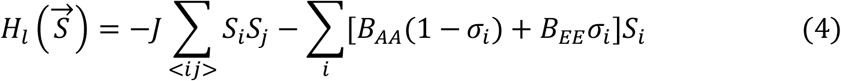

where the subunit arrangement (l) is specified by σ_*i*_(*i* = 1,2,…,6) that represents the identity of the *i*’th subunit in the hexamer ring: σ_*i*_ = 0 *or* 1 if *site* – *i* is AA or EE. The three parameters(*J*, *B_AA_*, *B_EE_*) are the same as those determined by using the previous experiments with pure KaiC-AA and KaiC-EE. Similar to Eq. (2), the probability of each hexamer conformational pattern *Pl*(*k*) can be calculated as:

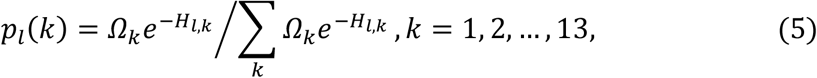

where *H_l,k_* is the energy for hexamer pattern-*k* with subunit arrangement-*l*. The overall distribution of the hexamer conformational patterns is obtained by the weighted average:

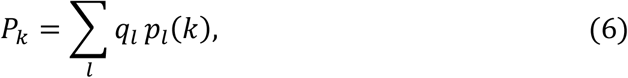

which depends on the distribution *q_l_*.

As a starting null hypothesis, we first considered two extreme scenarios for mixing: one is the fully-mixed scenario, *i.e*., each monomer in the ring has an equal probability of being AA or EE; another is the no-mixing scenario, *i.e*., only pure hexamers (all EE or all AA) exist. We found that neither of these extreme scenarios agrees with the data: *R*^2^ between the mixed experimental data and model prediction is 0.68 for no-mixing scenario (Fig. 4B) and 0.79 for fully-mixed scenario (Fig. 4C), see Supplementary Fig. 3 for the predicted distributions based on these scenarios. Our results thus rule out these two extreme scenarios.

Without detailed knowledge of the mixing process, we next treated *q_l_* as fitting parameters, and obtained their values by fitting the theoretical predicted *P_k_* (from Eqs. 5&6) to experimental observation (*P*_*k*(*ex*)_) subject to the constraints: 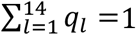, 0 ≤ *q_l_* ≤ 1; and the overall approximately equal percentage of KaiC-EE and KaiC-AA monomers over all hexamers (see Supplementary Note 2 for details). As shown in Fig. 4D, the model results are in good agreement with the experiments: *R*^2^ between the actual experimental data and model prediction data became 0.89. This agreement with experiments depends on the KaiC cooperativity. In the absence of the KaiC-KaiC interaction (J = 0), the agreement with experiments is poorer (*R*^2^ = 0.79) even when we allow *q_l_* to vary (see Supplementary Fig. 4A). Furthermore, the resulting distribution *q_l_* for *J* = 0 is almost the same as the no-mixing extreme case with a very small fraction of mixed hexamers (see Supplementary Fig. 5A), which is clearly inconsistent with our results for pure hexamers that show strong cooperativity. The better fit to data with a finite cooperativity *J* is robust when the weight constant (λ) in our optimization algorithm takes on different values (Supplementary Figs. 4 and 5B). Quantitatively, the fit of the Ising model to the mixed hexamer data is not as good as that in the pure hexamer case, which may indicate limitations of the two-state approximation; interactions between subunits may depend on their phosphorylation state in addition to their A-loop conformations (see Discussion and Supplementary Note 2 that incorporates this dependence).

We also examined the normalized correlation function 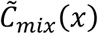, as shown in Fig.4E. It’s clear that 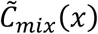 is non-zero for *x* ≥ 1, albeit smaller than that in the pure hexamer case. With the same fitting procedure for the correlation function as in the pure hexamer case, we obtained an effective coupling constant *J_mix_* = 0.086, which is about a half of that for pure hexamers. The statistically significant correlation thus confirms the existence of subunit-subunit cooperativity in mixed hexamers. However, the strength of cooperativity is weaker.

## Discussion

In this work, we report two distinct functional conformational states (exposed and buried) for each subunit that can coexist in a KaiC hexamer. Statistical analysis of single particle data suggests that there is a dynamic equilibrium between the two conformational states with the highly phosphorylated (dephosphorylated) KaiC prefers the buried (exposed) states, respectively. Characterization of the spatial arrangements of the exposed and buried states along with theoretical modeling reveal that there is substantial cooperativity in the hexamer that favors the neighboring KaiC monomers to be in the same conformational state. The day-night switch in Kai system is crucial in maintaining circadian rhythms, related to a general requirement for nonlinear response in the feedback loops that support chemical oscillators^34,35,39,50,51^. The cooperative interactions between neighboring KaiC subunits as identified in this study can make the transition from buried state to exposed state sharper, described by a higher Hill coefficient (Extended Data Fig. 10), and thus may provide a possible mechanism for controlling the day-night switch. The picture that emerges from our analysis is that interactions between the C-terminal regions within the KaiC hexamer form an intrinsically cooperative switch. The role of changing phosphorylation is then to shift the midpoint of the switch, ultimately actuating KaiB binding and closely the negative feedback loop to initiate the next cycle. We looked for structural differences in the CI domain and in the B-loop, but didn’t see any change up to the resolution of the structures. One possible scenario is that phosphorylation allows the CI domain to dynamically access conformations that are ultimately stabilized by KaiB binding but that are otherwise rare.

Our analysis of the mixed sample experiments suggests that the mixing of monomers with different phosphorylation levels may not be random. The enrichment factor α (the ratio of optimized *q_l_* values when λ = 0.1 (Supplementary Fig. 5B) to that under fully-mixed scenario (Supplementary Table 1)) indicates that there is a higher probability for two neighboring subunits to have different phosphorylation levels (Supplementary Fig. 5C). This possible preferential mixing phenomenon is worth further investigation given that KaiC hexamers constantly disassemble and reassemble during circadian oscillation through monomer shuffling ^27,52,53^, which is thought to be involved in synchronization^54–56^. In addition, we found that the effective coupling constant *J_mix_* in the Ising model for the mixed KaiC case is weaker than that in the pure hexamer case, which provide an alternative explanation for the reduced cooperativity due to a weaker coupling between heterogeneous KaiC monomers. Indeed, the microscopic origin of the nearest neighbor KaiC interaction that leads to cooperativity, which is key for coherent oscillations, is an important open question for future studies.

## Methods

### 1. Protein expression and purification

KaiC-AA and KaiC-EE were expressed and purified as described previously^39^. Briefly, KaiC mutants were expressed recombinantly in *E. coli* with N-terminal His_6_ tags. Clarified lysate was purified using Ni affinity chromatography (Histrap, Cytiva), fractions containing KaiC were pooled, and the expression tags were cleaved overnight. The resulting material was further purified using size exclusion chromatography (HiPrep S300, Cytiva) and fractions corresponding to the molecular weight of KaiC hexamers were selected.

### 2. Cryo-EM imaging and data collection

To remove the glycerol, KaiC proteins were applied to Zeba Micro Spin Desalting Columns (7K, Thermo Fisher), exchanging the buffer to running buffer (20 mM Tris-HCl (pH 8.0), 150mM NaCl, 5 mM MgCl2, 0.5 mM EDTA, 1 mM ATP). Then KaiC-AA (or KaiC-EE) was diluted to 0.35 ug/ul with running buffer and incubated at 30 °C for 6 hours to ensure that KaiC equilibrates to a functionally relevant state. Cryo-EM grids were prepared with FEI Vitrobot Mark IV. QUANTIFOIL grids (R2/1, 300 Mesh) were glow-discharged before a 3.5-μl drop of 0.35 ug/ul KaiC-AA (or KaiC-EE) solution was applied to the grids in an environmentally controlled chamber with 100% humidity and 4 °C temperature. After one blot force, 1 s blot time, the grid was plunged into liquid ethane and then was transferred to liquid nitrogen. The cryo-EM data was collected on a FEI Titan Krios G2 microscope connected to Gatan K2 Summit direct electron detector in a super-resolution counting mode, using SerialEM^57^ semi-automatically. Coma-free alignment was manually optimized and parallel illumination was verified before data collection. A total exposure time of 10 s with 250 ms per frame resulted in a 40-frame movie per exposure with an accumulated dose of ~50 electrons/Å^2^ (Extended Data Table 1). The calibrated physical pixel size and the super-resolution pixel size are 1.37 Å and 0.685 Å, respectively. Raw data were saved at the pixel size of 0.685 Å. A total of 5,125 movies of KaiC-AA and 3,530 movies of KaiC-EE were collected.

### 3. Preparation of mixed hexamer sample

The mixed sample was made following the method of Nishiwaki *et al^21^* to obtain KaiC-AA and KaiC-EE monomers. Protein concentrations were quantified by BCA Protein Assay Kit to prepare a 1:1 molar ratio of KaiC-AA to KaiC-EE. KaiC-AA (KaiC-EE) was buffer exchanged twice with Zeba Micro Spin Desalting Columns (7K, Thermo Fisher) into a buffer where ATP was replaced with 0.5 mM ADP, incubated at 4 °C for about 24 hours to disrupt hexamer structure. Then monomerized KaiC-AA and KaiC-EE were mixed for re-hexamerization by buffer exchanging into reaction buffer with 5 mM ATP^39^. Total KaiC concentration was quantified after re-hexamerization by BCA Protein Assay Kit, then diluted to 0.35 ug/ul with running buffer and incubated at 30 °C for 6 hours. Cryo-EM grid preparations and data collection procedures are the same as previously described.

### 4. Cryo-EM data processing

All frames of raw movies were aligned and averaged with the MotionCor2 program^58^ at a super-resolution pixel size of 0.685 Å. Each drift-corrected micrograph was used for the determination of the micrograph CTF parameters with program Gctf ^59^. We picked 1,592,573 particles of the KaiC-AA, 934,373 particles of the KaiC-EE, 693,666 particles of the mixed hexamer sample using the program EMAN2^60^. Reference-free 2D classification and 3D classification were carried out with two-fold binned data with a pixel size of 1.37 Å in both RELION^44^-^45^ and ROME^61^. Focused 3D classification, which we used in the later stage of data processing, and high-resolution refinement were mainly conducted with RELION^44,45^. A substantial part of the data processing, mostly 2D/3D classification, was performed with clusters supported by High Performance Computing Platform in PKU.

There were 140,475 particles of KaiC-AA, 181,326 particles of KaiC-EE, 175,284 particles of the mixed hexamer sample in the dataset chosen for the following steps of analysis. The final refinement was done using data with a pixel size of 1.37 Å that were binned by two-fold from the raw data in the super-counting mode. Based on the in-plane shift and Euler angle of each particle from the last iteration of refinement, we reconstructed the two half-maps of each structure using raw single particle images at the super-resolution mode with a pixel size of 0.685 Å, which resulted in reconstructions for the KaiC-AA, KaiC-EE, and the mixed sample with overall resolutions of 3.3 Å, 3.3 Å and 3.8 Å, respectively, measured by gold-standard FSC at 0.143-cutoff. All density maps were sharpened by applying a negative *B*-factor −100 manually. Local resolution variations were further estimated using ResMap^62^.

### 5. Atomic model building and refinement

To build the initial atomic models of KaiC-AA and KaiC-EE, we used a previously published KaiC structure^47^ and then manually improved the main-chain and side-chain fitting in Coot^63^ to generate the starting coordinate files. To fit the KaiC-AA and KaiC-EE atomic models to the corresponding reconstructed density maps, we first conducted rigid body fitting of the segments of the model in Chimera^64^, after which the fitting of atomic models with density maps were improved manually in Coot. Atomic models of *S_Ex1_, S_Ex2_, S_Bu1_, S_Bu2_, S_Bu3_* were fitted in Coot manually starting from the KaiC-EE structure. Finally, atomic models were all subjected to the real-space refinement program in Phenix ^65^, see Extended Data Table 1 for validation statistics.

### 6. Structural analysis and visualization

Structural comparison was conducted in Pymol^66^, Chimera^64^ and ChimeraX^67^. All figures of the structures were plotted using Pymol^66^, Chimera^64^ and ChimeraX^67^.

### 7. Six-fold pseudo-symmetry and classification results by RELION

We used 140,475 particles of KaiC-AA (Extended Data Fig. 4), 371,557 particles of KaiC-EE (two remaining classes were added, which also resulted in good-quality refinement with overall resolutions of 3.8 Å) (Extended Data Fig. 6), and 175,284 particles of the mixed sample (Extended Data Fig. 8D) to perform C6 pseudo-symmetry expansions in RELION^44,45^. That is rotating each particle by 60°, 120°, 180°, 240°, 300° to get the symmetric copies and expanding the particle set. In this way, the whole data set was expanded 6 times. We used focused 3D classification (with CII domain masked) to further classify particles into two different conformational states (exposed state and buried state) using RELION^44,45^, see Extended Data Fig. 5,7 and 9 for the criterion of these states.

### 8. Criteria for distinguishing the exposed (Ex) state and buried (Bu) state

For each dataset (EE, AA, and mixed), we used the average 3D volumes of those clusters with the most well-established buried structure within the A-loop area (cluster 1-4 in Extended Data Fig. 5 for KaiC-EE; clusters 1-3, 1-7, 1-8, 1-10, 1-11 and 1-13 in Extended Data Fig. 7 for KaiC-AA; clusters 1-3, 1-6, 1-15 in Extended Data Fig. 9 for mixed KaiC) to create a black-and-white mask (digital map) by using a density threshold (0.025 for KaiC-AA, 0.2 for KaiC-EE, 0.1 for mixed KaiC) in RELION^44,45^. We then extend the white volume by one (or two) pixels in all directions to obtain mask1 (or mask2). The overlap intensities (or the integral density values) of the n-th 3D volume with mask1 (or mask2) is defined as *I*_1,*n*_ = ∑*iD_i,n_M*_1,*i*_ (*or I*_2,*n*_ = ∑_i_*D_i,n_M*_2,*i*_), *D_i,n_* is the density value of the n-th 3D volume at the i-th pixel point and *M*_1,*i*_ (*M*_2,*i*_) is a binary number that is 1 inside the white volume of mask 1 (mask 2) and 0 in the black volume of mask 1 (mask 2). The 3D volumes with larger integral density values of *I*_1,*n*_ and *I*_2,*n*_ are more likely to be in the buried (Bu) state, and those with smaller values of *I*_1,*n*_ and *I*_2,*n*_ are more likely to be in the extended (Ex) state. To increase the statistical confidence of the classification of the two states (Ex and Bu), we consider the 3D volumes with intermediate values of *I*_1,*n*_ and *I*_2,*n*_ to have an undefined (Un) conformational state. For details, see Extended Data Figs. 5A, 7A and 9A. To check the robustness and consistency of the classification of the buried (Bu) and the exposed (Ex) state, and more importantly, to independently judge the state of those 3D volumes at the boundaries between Un and either the Bu or the Ex states, each 3D volume is shown (only within the region corresponding to mask2) at high density threshold (4σ) and low density threshold (2σ), where σ is the standard deviation of the 3D volume (density) within the hexamer region, see Extended Data Figs. 5B, 7B and 9B. These 3D volumes are inspected to help determine their conformational states, especially at the boundary between the Bu state and the Un state where the determination is difficult by using the overlap intensities alone.

## Data availability

The three-dimensional cryo-EM density maps of KaiC-AA and KaiC-EE are deposited into the Electron Microscopy Data Bank (EMDB) KaiC-AA: EMD-32952, KaiC-EE: EMD-32953. The corresponding atomic coordinates are deposited in the Protein Data Bank (PDB) KaiC-AA: 7X1Y, KaiC-EE: 7X1Z.

## Acknowledgments

The authors thank Y. D. Mao, Y. N. Zhu, B. Liu, S. W. Zhang, S. T. Zou, H. Liang, Y. H. Wang, K. Y. Wang, D. Y. Yin and W. L. Wang for constructive discussions; Y. Ma, X. Li, X. Pei for technical supports. The cryo-EM data were collected from the Electron Microscopy Laboratory and cryo-EM Platform at Peking University. Data processing was performed on the Weiming No.1 and Life Science No. 1 High Performance Computing Platform in Peking University. The work by D.Z. is supported by the China Postdoctoral Science Foundation (2020M680180).

M. J. Rust is supported by an HHMI-Simons Faculty Scholar award and NIH R01 GM107369. The work by Y. T. is supported by a NIH grant (R35GM131734). The work by Q.O. is supported by the National Natural Science Foundation of China (12090054).

## Author contributions

L.H. purified proteins; X.H. prepared samples for imaging and collected data; X.H., D.Y. processed data and refined the maps; X.H., D.Z. did the simulations and analyzed the data; Z.W., T.Y. contributed to simulations and data analysis; M.R., Y.T., Q.O. initiated the project, developed the model and analyzed the data; all wrote the paper.

## Notes

### Competing Interest Statement

The authors have declared no competing interest.

### Summary of Updates

Several minor errors due to format conversion revised.

