## Extended Data Table 1 and Figures 1 to 10 for "A cooperative switch within the KaiC hexamer revealed by cryo-EM"

**Extended Data Table 1. Cryo-EM data collection, refinement and validation statistics.**

| | KaiC-AA | KaiC-EE |
|---|---|---|
| **Data collection and processing** | | |
| Magnification | 105,000 | 105,000 |
| Voltage (kV) | 300 | 300 |
| Electron exposure (e–/Å$^2$) | 50 | 50 |
| Defocus range (μm) | -0.4 to -5.0 | -0.4 to -5.0 |
| ixel size (Å) | 0.685 | 0.685 |
| Symmetry imposed | C6 | C6 |
| Initial particle images (no.) | 1,592,573 | 934,373 |
| Final particle images (no.) | 140,475 | 181,326 |
| Map resolution (Å) | 3.3 | 3.3 |
| FSC threshold | 0.143 | 0.143 |
| **Refinement** | | |
| Map sharpening *B* factor (Å$^2$) | -100 | -100 |
| **Model composition** | | |
| Non-hydrogen atoms | 22,026 | 23,298 |
| Protein residues | 2748 | 2,904 |
| Ligands | 24 | 24 |
| **R.m.s. deviations** | | |
| Bond lengths (Å) | 0.006 | 0.003 |
| Bond angles (°) | 0.852 | 0.753 |
| **Validation** | | |
| MolProbity score | 1.65 | 1.67 |
| Clashscore | 3.60 | 4.99 |
| Rotamers outliers (%) | 0.00 | 0.04 |
| **Ramachandran plot** | | |
| Favored (%) | 91.72 | 93.84 |
| Allowed (%) | 8.28 | 6.16 |
| Outliers (%) | 0.00 | 0.00 |

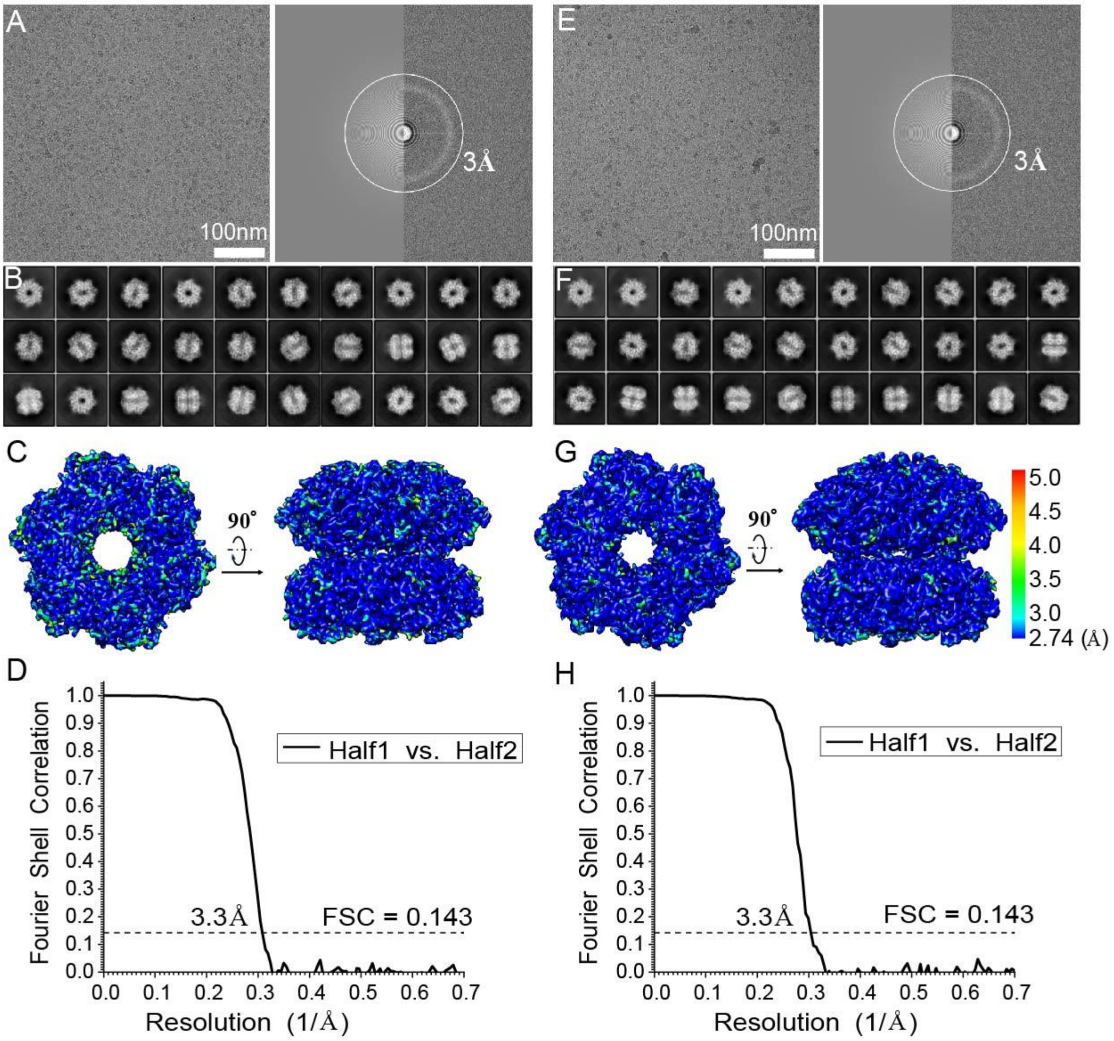

**Extended Data Fig. 1. Cryo-EM structure determination of KaiC-AA and KaiC-EE.** On the left is a typical cryo-EM micrograph of KaiC-AA **(A)** and of KaiC-EE **(E)**, taken with a FEI Titan Krios G2 microscope equipped with the post-column Gatan BioQuantum energy filter connected to Gatan K2 Summit direct electron detector. Scale bar, 100 nm. On the right is the power spectrum evaluation corresponds to the left micrograph. (B) and (F) give gallery of unsupervised 2D class averages of KaiC-AA **(B)** and of KaiC-EE **(F)**. (C) and (G) provide local resolution estimation calculated by ResMap for KaiC-AA **(C)** and for KaiC-EE **(G)**. (D) and (H) show Fourier shell correlation (FSC) for two independently refined halves of KaiC-AA **(D)** and of KaiC-EE **(H)**. During the refinement we imposed C6 symmetry.

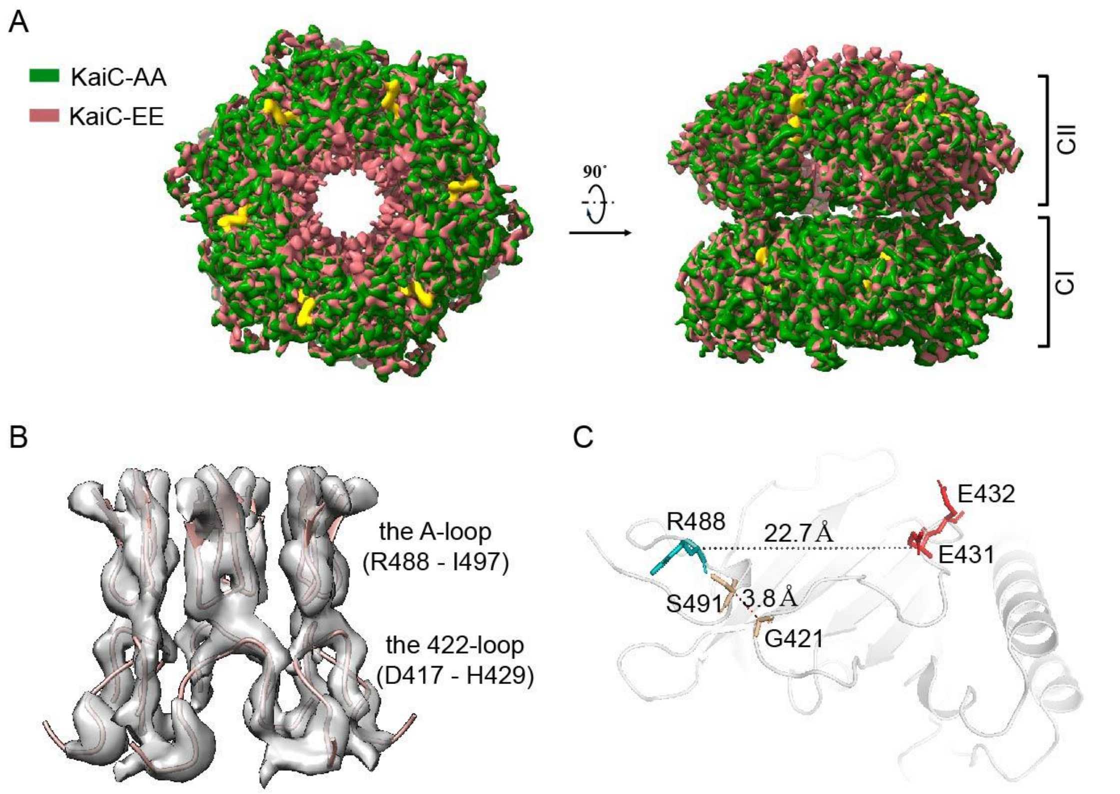

**Extended Data Fig. 2. Comparison of KaiC-AA and KaiC-EE cryo-EM densities. (A)** Superimposition of KaiC-AA density (colored in dark green) and KaiC-EE density (colored in red). **(B)** The difference map between KaiC-EE and KaiC-AA cryo-EM density is rendered as a transparent surface, superimposed with the cartoon representation of the atomic model of the A-loop (residue 488-497) and the 422-loop (residue 417-429). **(C)** A close-up view of this interacting-loops region in KaiC-EE CII domain (subunit A). Side chains of G421, E431, E432, R488, S491 and distance between G421 and S491 are labeled.

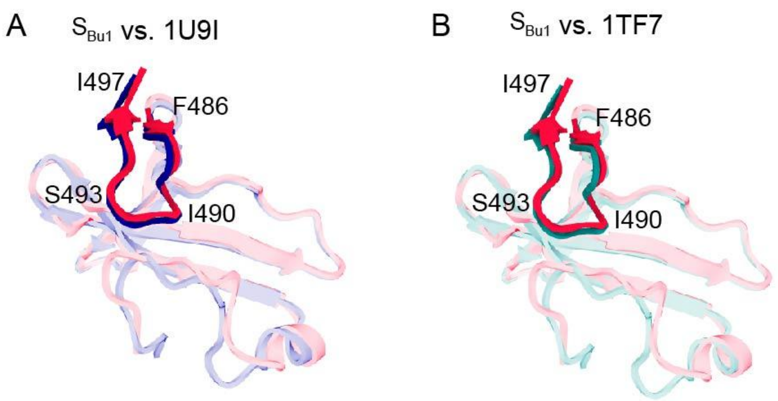

**Extended Data Fig. 3 (A)** and **(B) The buried state ($S_{Bu1}$) superimposed with previously reported crystal structures in PDB data bank (with 1U9I, 1TF7 colored in dark blue and teal, respectively).** The A-loop is shown as cartoons without transparency.

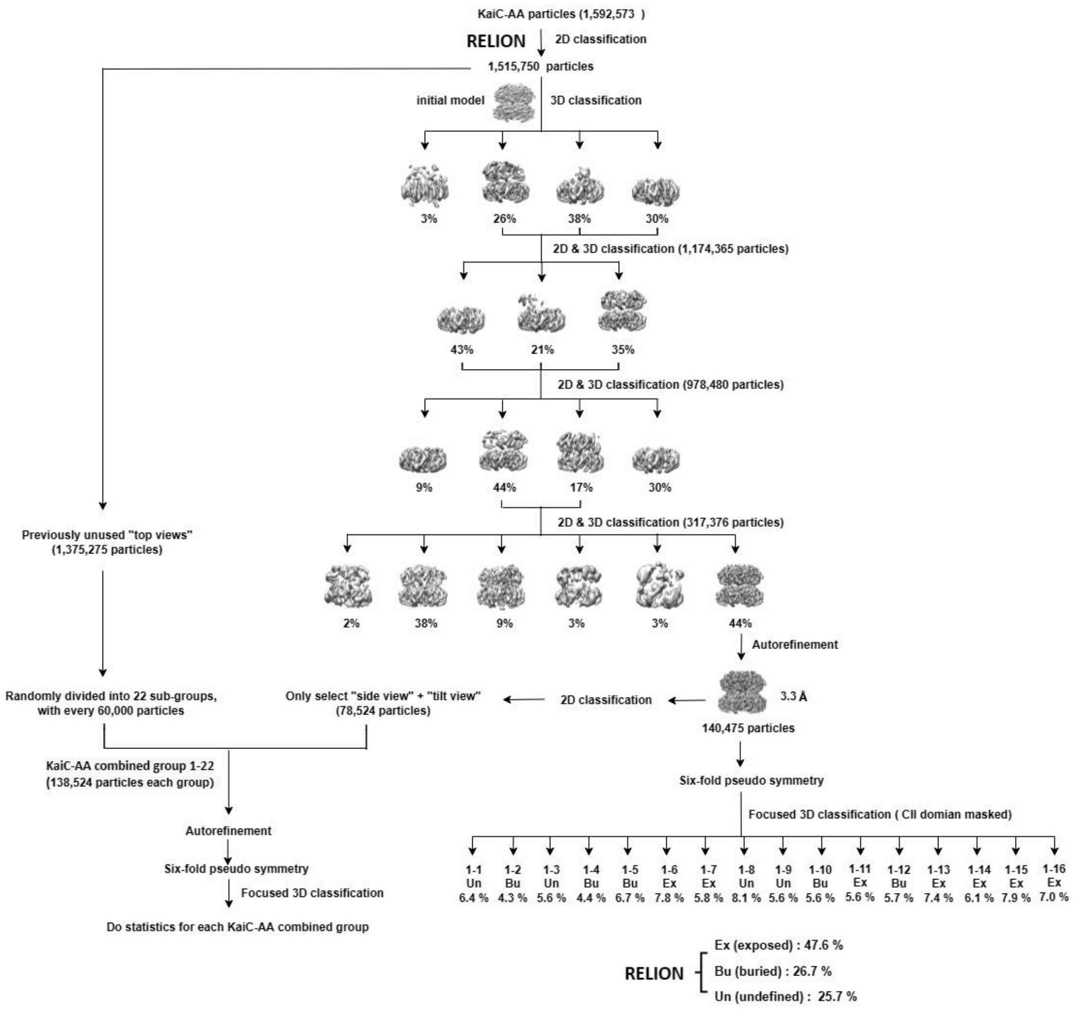

**Extended Data Fig. 4. Data processing flow chart of KaiC-AA data set.** For the 3D classification results of the high quality well-defined structure (consist of 140,475 particles), there is 47.6% of particles belong to the exposed state and 26.7% of particles belong to the buried state, the rest 25.7% particles remain unclassified. With *x*–*y* shift and angular information, we can put these exposed/buried subunits back into hexamers, and then carry out statistical analysis of the hexamer conformational patterns shown in Fig. 3A in the main text.

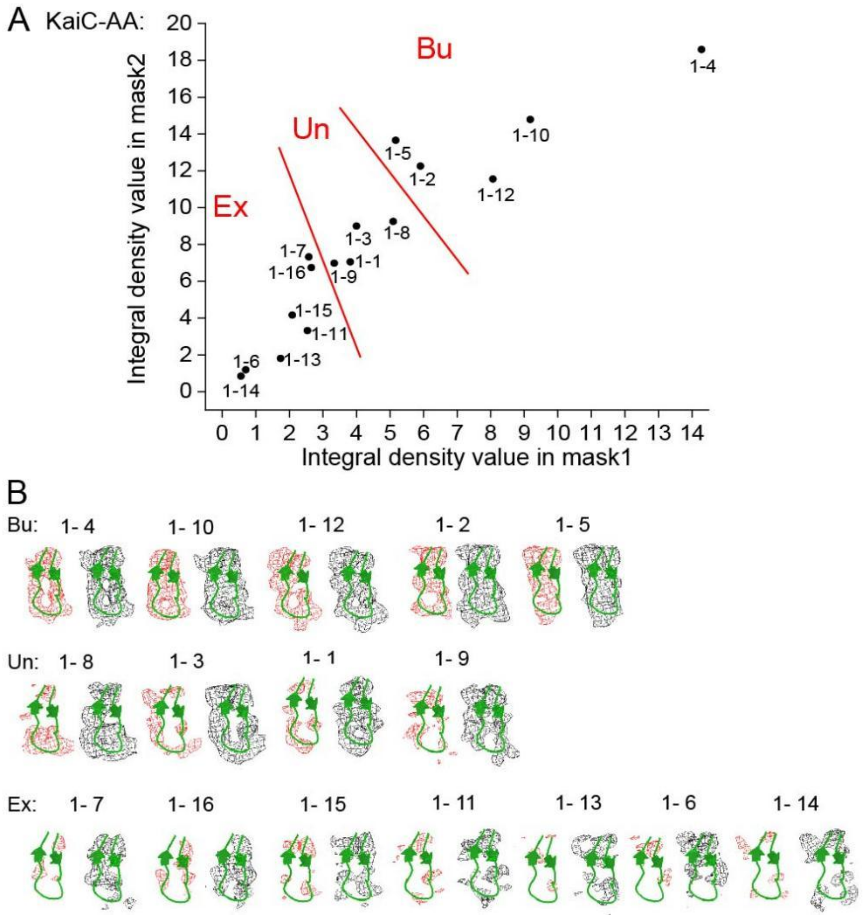

**Extended Data Fig. 5. Criteria for distinguishing the exposed (Ex) state and buried (Bu) state in KaiC-AA.** (A) The overlap intensities (Integral density values) of 3D volumes classified by RELION (Extended Data Fig. 4) with the two masks that are used to characterize the buried state (Bu). (B) All 3D volumes (within the region corresponding to mask2) are shown at a high density threshold ($4\sigma$, red mesh, left) and a low density threshold ($2\sigma$, black mesh, right). The first row shows the structures that are classified as the buried (Bu) state. All structures in the first row agree with atomic model in all parts of the A-loop area at both the high and the low thresholds. The second row shows the undefined (Un) states, with density profiles that agree with the atomic model of the Bu state at the low threshold but not at the high threshold. The third row shows exposed (Ex) states, with the density profiles that disagree with the atomic model of the Bu state in the A-loop area at both the high and the low thresholds.

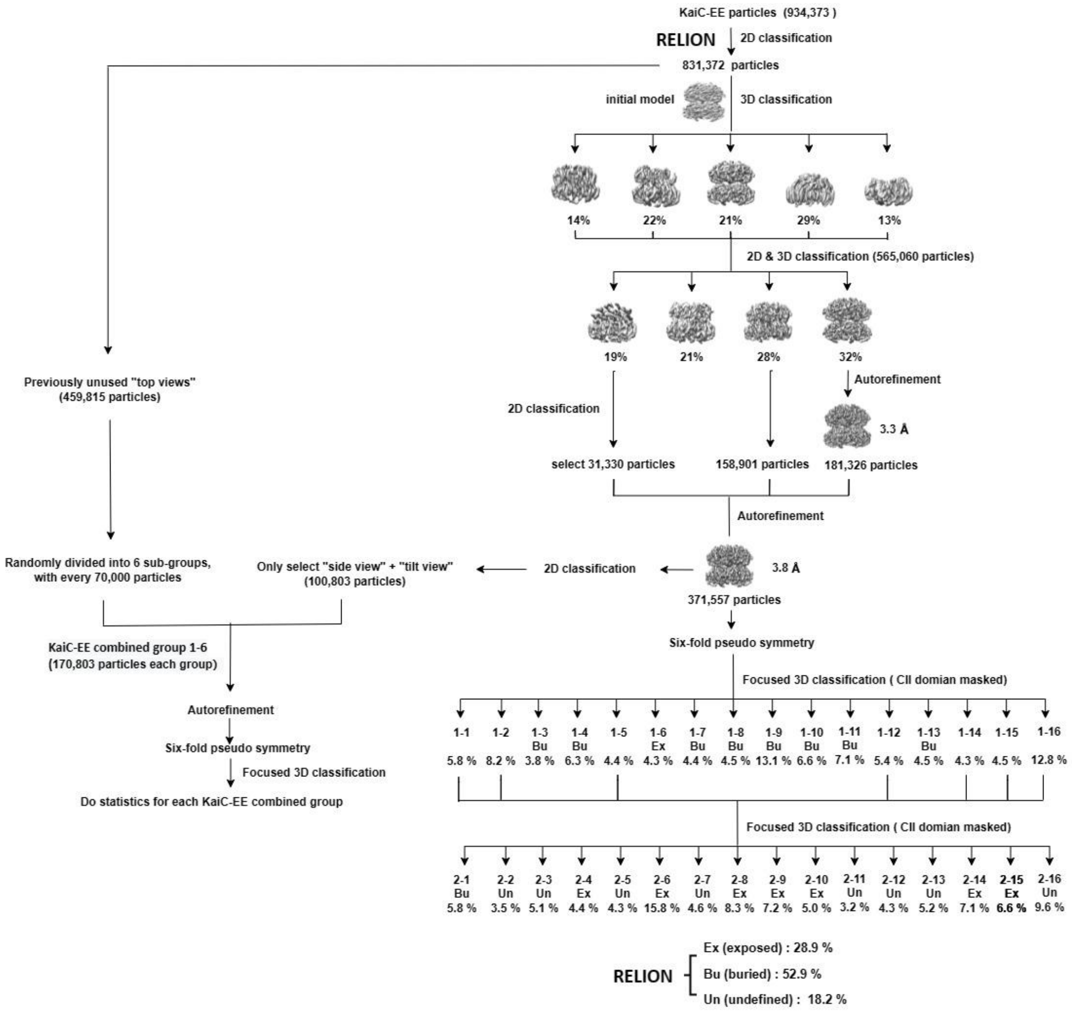

**Extended Data Fig. 6. Data processing flow chart of KaiC-EE data set.** For the 3D classification results of the high quality well-defined structure (consist of 371,557 particles), there is 28.9% of particles belong to the exposed state and 52.9% of particles belong to the buried state, the rest 18.2% particles remain unclassified. With *x*–*y* shift and angular information, we can put these exposed/buried subunits back into hexamers, and then carry out statistical analysis of the hexamer conformational patterns shown in Fig. 3B in the main text.

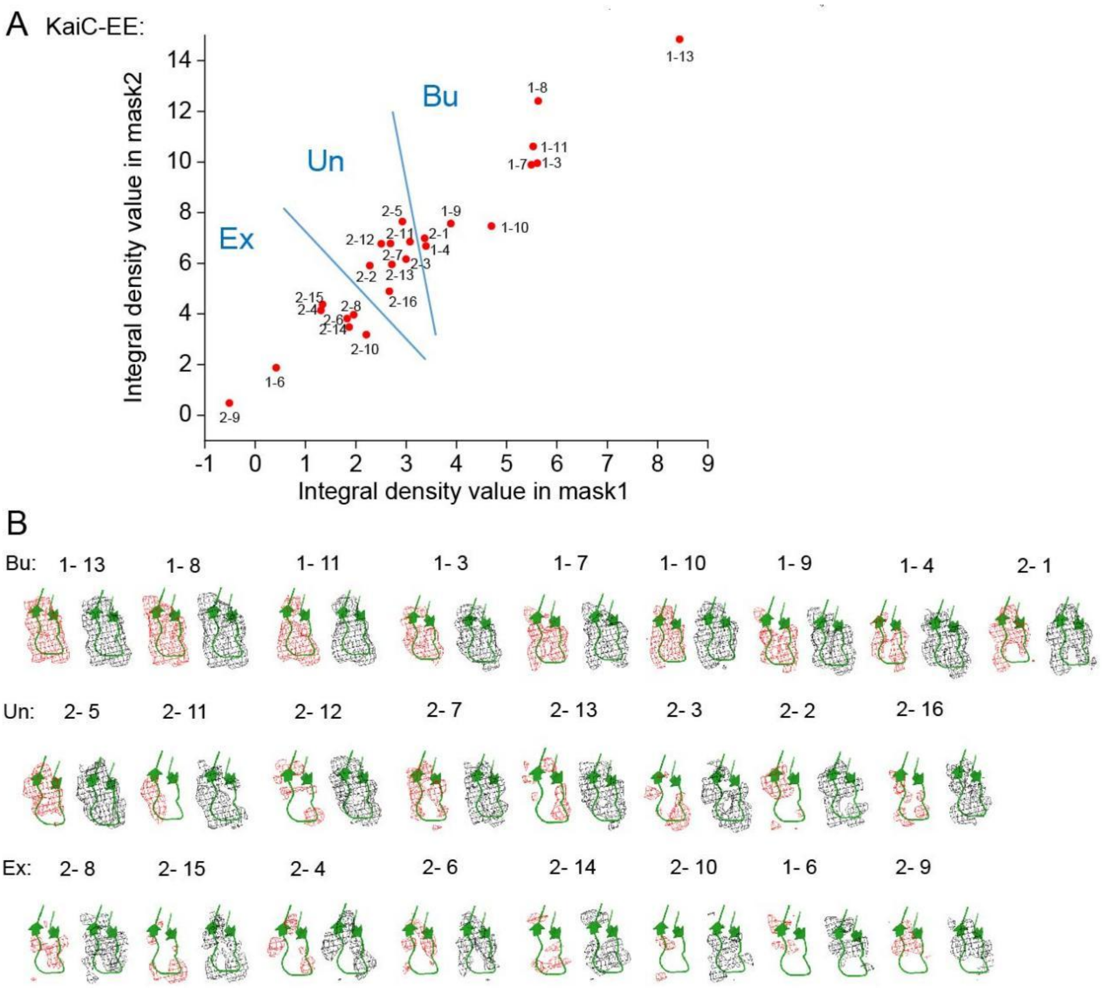

**Extended Data Fig. 7. Criteria for distinguishing the exposed (Ex) state and buried (Bu) state in KaiC-EE.** (A) The overlap intensities (Integral density values) of 3D volumes classified by RELION (Extended Data Fig. 4) with the two masks that are used to characterize the buried state (Bu). (B) All 3D volumes (within the region corresponding to mask2) are shown at a high density threshold ($4\sigma$, red mesh, left) and a low density threshold ($2\sigma$, black mesh, right). The first row shows the buried (Bu) states, the second row shows undefined (Un) states, the third row shows exposed (Ex) states. Note that the 3D volume 2-5 has a poor resolution and weak density at the bottom at the $4\sigma$ density threshold, and the 3D volume 2-11 only has density on the left-side at the $4\sigma$ density threshold. So both 2-5 and 2-11, which are at the boundary of the Un states and the Bu states, are classified as undefined (Un). However, even including 2-5 as a buried state does not affect the final results.

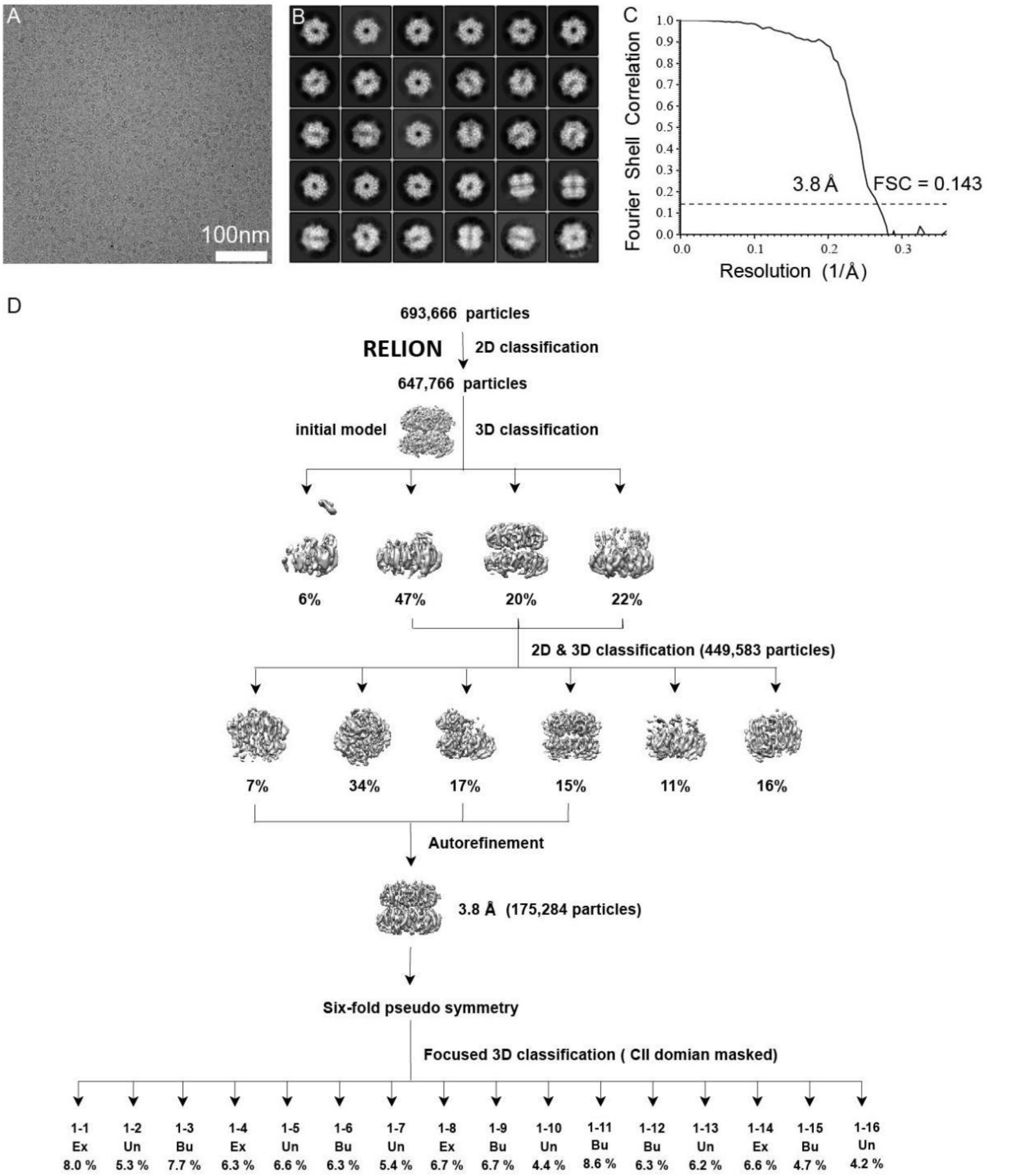

**Extended Data Fig. 8. Cryo-EM structure determination and data processing flow chart of the mixed sample data set. (A)** Typical cryo-EM micrograph of the mixed sample data set, taken with a FEI Titan Krios G2 microscope equipped with the post-column Gatan BioQuantum energy filter connected to Gatan K2 Summit direct electron detector. Scale bar, 100 nm. **(B)** Gallery of unsupervised 2D class averages of the mixed sample data set. **(C)** Fourier shell correlation (FSC) for two independently refined halves of the mixed sample data set. During the refinement we imposed C6 symmetry. **(D)** Data processing flow chart of the mixed sample data set.

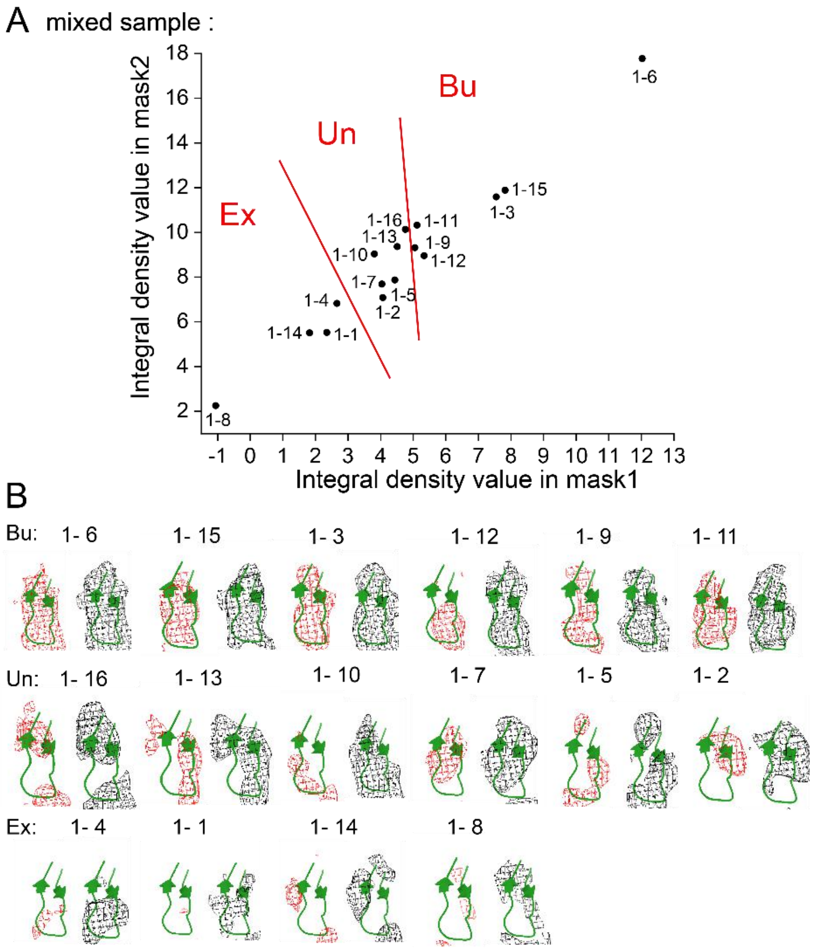

**Extended Data Fig. 9. Criteria for distinguishing the exposed (Ex) state and buried (Bu) state in mixed sample.** ((A) The overlap intensities (Integral density values) of 3D volumes classified by RELION (Extended Data Fig. 4) with the two masks that are used to characterize the buried state (Bu). (B) All 3D volumes (within the region corresponding to mask2) are shown at a high density threshold ($4\sigma$, red mesh, left) and a low density threshold ($2\sigma$, black mesh, right). The first row shows the buried (Bu) states, the second row shows the undefined (Un) states, the third row shows the exposed (Ex) states. Note that the 3D volume 1-16 and 1-13, which at the boundary between the Un state and the Bu state, have poor resolution and weak densities at the $4\sigma$ density threshold, so they are classified as undefined (Un).

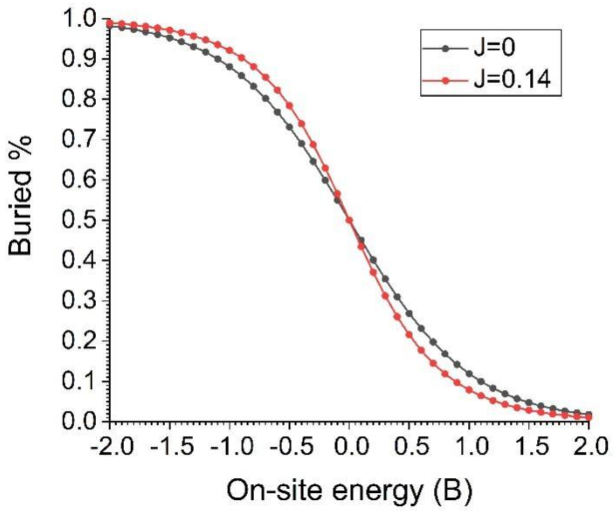

**Extended Data Fig. 10. The percentage of buried state as we vary the on-site energy ($B$) for without coupling ($J = 0$) and with coupling ($J = 0.14$).**
