## Supplementary Information for "A cooperative switch within the KaiC hexamer revealed by cryo-EM"

### Supplementary Note 1: Statistics of pattern distribution in pure hexamers

In this note, we briefly explain some critical points and introduce additional fitting methods and results mentioned in the main text, regarding the statistics of hexamer pattern distribution in pure hexamers.

**The conformational patterns of hexamer.** As mentioned in the main text, each monomer could be either in the exposed (Ex) state or buried (Bu) state, so there should be $2^6 = 64$ possible hexamer configurations in total. Because all the monomers are regarded identical, the configurations that only differ in a rotation operation can be grouped into the same hexamer conformational pattern, as is shown in Supplementary Fig. 1, and the number of configurations in the group is the pattern’s degeneracy. Furthermore, for the two patterns that only differ in chirality (pattern-(7) and pattern-(8) in Supplementary Fig.1A), we found that their numbers (in KaiC-AA, pattern-(7) has 1,292 particles pattern-(8) has 1,399 particles; in KaiC-EE, pattern-(7) has 6,306 particles pattern-(8) has 6,248 particles) are quite close. Hence, we ignore this chiral effect and regard the two patterns the same. So, there are 13 patterns in total, whose degeneracies are listed in Supplementary Fig. 1B.

**Processing of previously unused particles.** To avoid bias in particle selection, we also carried out statistical analysis by including a large number of particles that were previously unused in the main text due to orientation preference (the previously unused “top views”, including 1,375,275 KaiC-AA particles and 459,815 KaiC-EE particles, see Extended Data Figs. 4 and 6). These particles were randomly divided into sub-groups with equal number of particles (22 groups for KaiC-AA and 6 groups for KaiC-EE). Then, each sub-group were combined with “side view” and “tilt view” particles selected from the 2D classification of the high quality well-defined structures (140,475 particles for KaiC-AA and 371,557 particles for KaiC-EE), respectively. These combined particles are numbered from “KaiC-AA combined group 1” to “KaiC-AA combined group 22” (or from “KaiC-EE combined group 1” to “KaiC-EE combined group 6”). Finally, we performed statistical analysis for each combined group following the same procedures described in the main text by using RELION (Extended Data Figs. 4 and 6). We show the statistical results of two combined groups for KaiC-AA and KaiC-EE respectively in Supplementary Fig. 2, which indicates that including these previously unused particles did not qualitatively change statistics of the hexamer patterns.

### Supplementary Note 2: Statistics of pattern distribution in mixed hexamers

In this note, we describe some details about conformational statistics for mixed hexamers.

**Conformational pattern distribution and monomer arrangement for mixed hexamers.**

In principle, the conformational pattern distribution for mixed hexamers should lie between the two pure cases. This is because the mixed hexamers can have 14 distinct monomer arrangements (see Supplementary Table 1), though the arrangement distribution $q_l$ is unknown. Two extreme scenarios to infer $q_l$ is assuming either monomers are: 1) fully mixed, i.e., AA and EE monomers appear in the ring with no preference (fully-mixed scenario), or 2) not mixed at all (the no-mixing scenario). In fully-mixed scenario, $q_l$ can be calculated analytically and listed in Supplementary Table 1. In no-mixing scenario, only the two pure arrangements have non-zero $q_l$. With the two assumptions of $q_l$, we can calculate the theoretical distribution of hexamer patterns in respective scenarios. These two theoretical distributions are shown in Supplementary Fig. 3, with the three experimental distribution (two pure cases and the mixing case) also presented for comparison. It appears that neither of the two extreme scenarios can well explain the experimental distribution of mixed hexamers.

**The fitting for subunit arrangement.** Generally, $q_l$ could be obtained from fitting. We fit the theoretical predicted $P_k$ (according to Eqs. 5 & 6 in main text) to experimental observation ($P_{k(ex)}$) by using the following least square method:

$$min_q\left(\sum_{k=1}^{13}\left(\sum_{l=1}^{14} q_l p_l(k) - P_{k\ (ex)}\right)^2 + \lambda\left(\sum_{l=1}^{14} q_l(1-E_l) - \sum_{l=1}^{14} q_l E_l\right)^2\right), \quad (1)$$

subject to the constraints: $\sum_{l=1}^{14} q_l = 1,\ \ 0 \le q_l \le 1$. The second term above is introduced to enforce the overall equal percentage of KaiC-EE and KaiC-AA monomers with $E_l$ the percentage of KaiC-EE monomer in the hexamer with arrangement-$l$, and λ is a weight constant. The fitting results $q_l$ and performance $R^2$ are in general insensitive to λ (Supplementary Figs. 4 & 5B). We choose a specific $\lambda = 0.1$ in the analysis in main text to enforce the difference between KaiC-AA and KaiC-EE percentage to be less than 10%.

From the fitting result of $q_l$ shown in Supplementary Fig. 5A, in the absence of coupling, the resulting distribution is almost the same as no-mixing scenario, which is inconsistent with our previous results and certainly not the case. Hence strong coupling is necessary.

A

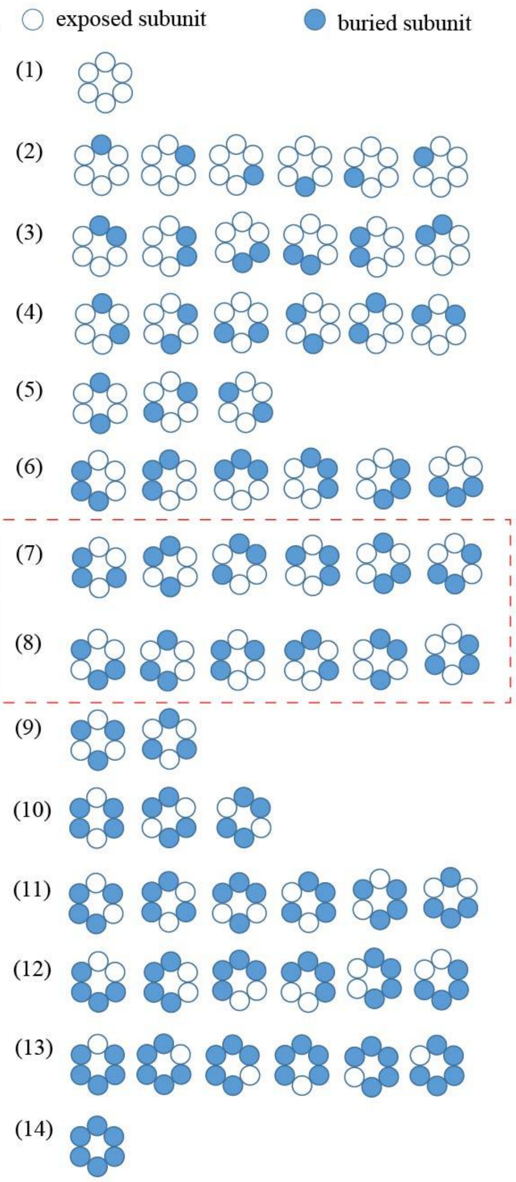


B

| pattern | the degeneracy $\Omega_k$ | $n_k$ |
|---|---|---|
| 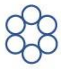 | $\Omega_1 = 1$ | $n_1 = 6$ |
| 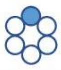 | $\Omega_2 = 6$ | $n_2 = 5$ |
| 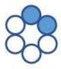 | $\Omega_3 = 6$ | $n_3 = 4$ |
| 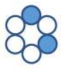 | $\Omega_4 = 6$ | $n_4 = 4$ |
| 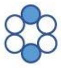 | $\Omega_5 = 3$ | $n_5 = 4$ |
| 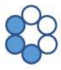 | $\Omega_6 = 6$ | $n_6 = 3$ |
| 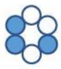 | $\Omega_7 = 12$ | $n_7 = 3$ |
| 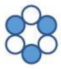 | $\Omega_8 = 2$ | $n_8 = 3$ |
| 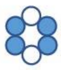 | $\Omega_9 = 3$ | $n_9 = 2$ |
| 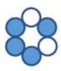 | $\Omega_{10} = 6$ | $n_{10} = 2$ |
| 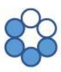 | $\Omega_{11} = 6$ | $n_{11} = 2$ |
| 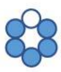 | $\Omega_{12} = 6$ | $n_{12} = 1$ |
| 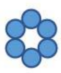 | $\Omega_{13} = 1$ | $n_{13} = 0$ |

**Supplementary Fig. 1. Considering the structure degeneracy, 64 configurations can be combined into 13 conformational patterns. (A)** These 64 configurations are grouped into 13 conformational patterns according to the same pattern. Note that pattern (7) and (8) are only different in chirality, and our experiment shows very little difference between their numbers. So these two patterns are combined into one. **(B)** The degeneracy $\Omega_k$ and the number of exposed state $n_k$ are listed for each hexamer pattern.

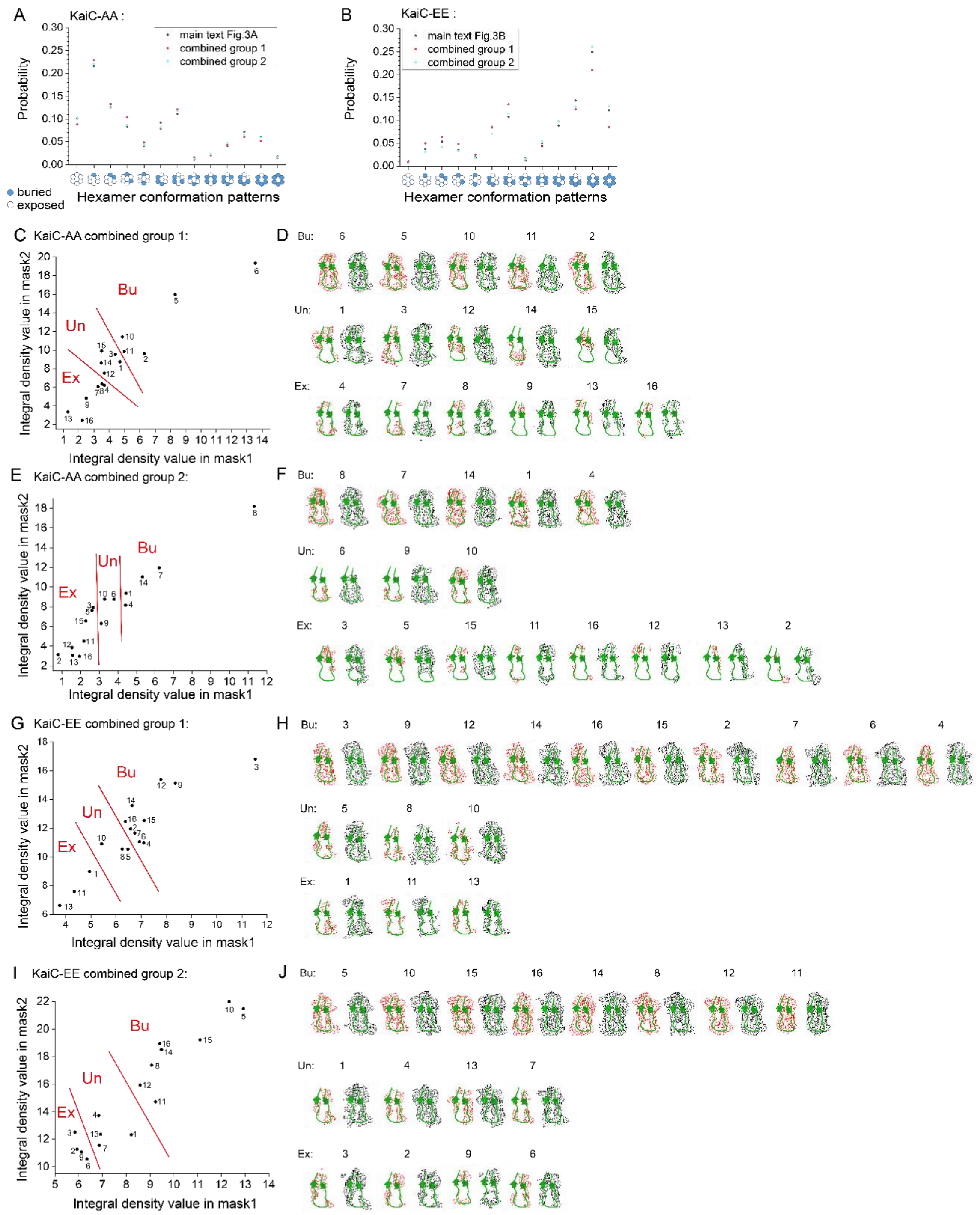

**Supplementary Fig. 2. Including previously unused particles due to orientation preference does not change the statistical results. (A)** Comparison of the hexamer conformational pattern distribution in main text Fig. 3A (24,240 rings) and that of KaiC-AA combined group 1 (20,133 rings) and combined group 2 (21,638 rings). **(B)** Comparison of the hexamer conformational pattern distribution in main text Fig. 3B (116,785 rings) and that of KaiC-EE combined group 1 (84,226 rings) and combined group 2 (13,536 rings)). Probability of each data set was normalized. **(C-J)** The overlap intensities (Integral density values) of 3D volumes classified by RELION with the two masks that are used to characterize the buried state (Bu) for KaiC-AA combined group 1 **(C)**, KaiC-AA combined group 2 **(E)**, KaiC-EE combined group 1 **(G)**, KaiC-EE combined group 2 **(I)**. All 3D volumes (within the region corresponding to mask2) are shown at a high density threshold ($4\sigma$, red mesh, left) and a low density threshold

($2\sigma$, black mesh, right) for KaiC-AA combined-group 1 **(D)**, KaiC-AA combined-group 2 **(F)**, KaiC-EE combined-group 1 **(H)**, KaiC-EE combined-group 2 **(J)**.

**Supplementary Table 1. Enumeration of subunit arrangements. The arrangements for the two extreme scenarios are also listed.**

| subunit arrangements | $q_l$ | fully-mixed | no-mixing |
|---|---|---|---|
| AAAAAA | $q_1$ | 0.0156 | 0.5 |
| AAAAAE | $q_2$ | 0.0938 | 0 |
| AAAAEE | $q_3$ | 0.0938 | 0 |
| AAAEAE | $q_4$ | 0.0938 | 0 |
| AAEAAE | $q_5$ | 0.0469 | 0 |
| AAAEEE | $q_6$ | 0.0938 | 0 |
| AAEAEE | $q_7$ | 0.0938 | 0 |
| AAEEAE | $q_8$ | 0.0938 | 0 |
| AEAEAE | $q_9$ | 0.0313 | 0 |
| AAEEEE | $q_{10}$ | 0.0938 | 0 |
| AEAEEE | $q_{11}$ | 0.0938 | 0 |
| AEEAEE | $q_{12}$ | 0.0469 | 0 |
| AEEEEE | $q_{13}$ | 0.0938 | 0 |
| EEEEEE | $q_{14}$ | 0.0156 | 0.5 |

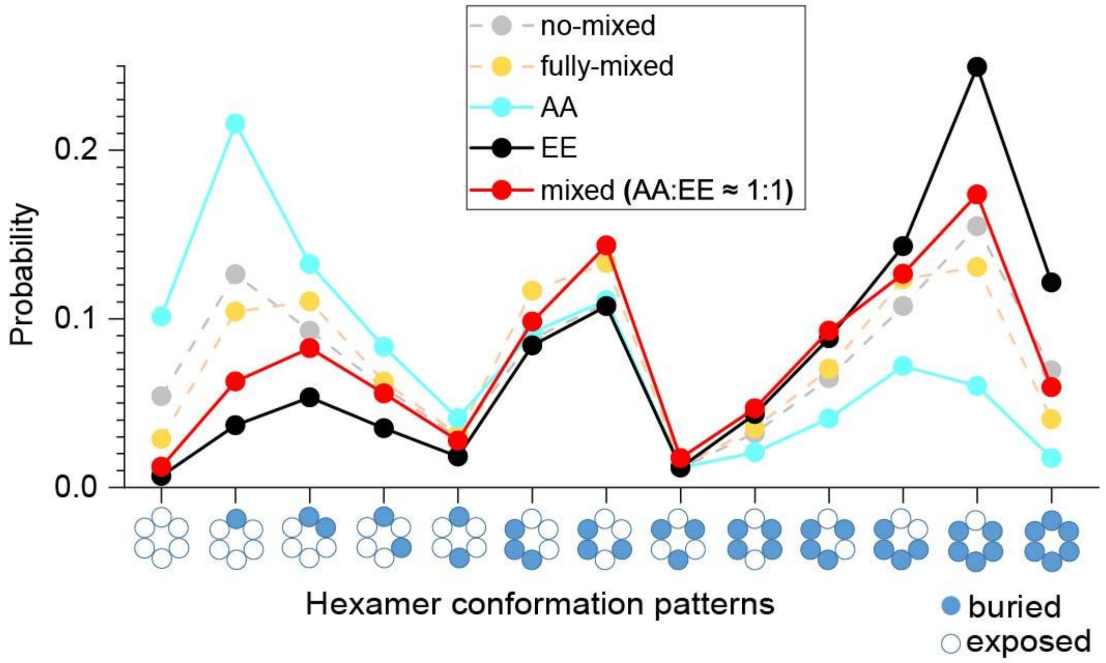


**Supplementary Fig. 3. Hexamer conformation pattern** distributions, predicted in two extreme scenarios (connected by dotted line) and measured in three experimental conditions (connected by solid line). Predicted distribution in no-mixing scenario and fully-mixed scenario are connected by gray dotted line and yellow dotted line, respectively. Experimental distribution of pure KaiC-AA, pure KaiC-EE and the mixed sample are connected by cyan solid line, black solid line and red solid line, respectively. The conformation distribution of mixed hexamers lies between two pure distributions, but can be explained neither by no-mixing assumption nor by fully-mixing assumption.

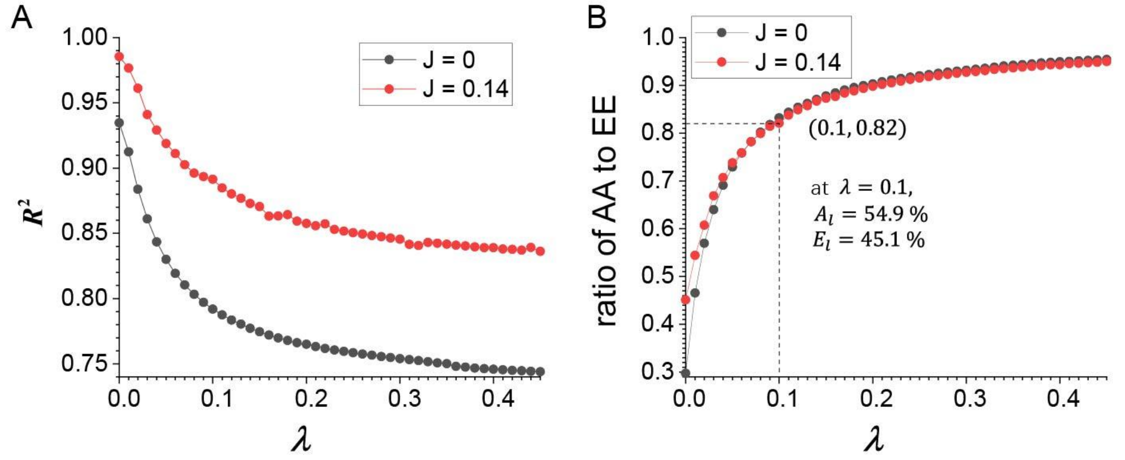


**Supplementary Fig. 4. (A)** $R^2$ versus λ when the coupling constant $\mathrm{J} = 0$ (black) or $\mathrm{J} = 0.14$ (red). **(B)** The ratio of AA monomer to EE monomer for the fitted arrangement distribution varies with λ when the coupling constant $\mathrm{J} = 0$ (black) or $\mathrm{J} = 0.14$ (red). The fitting performance is quite robust when λ varies.

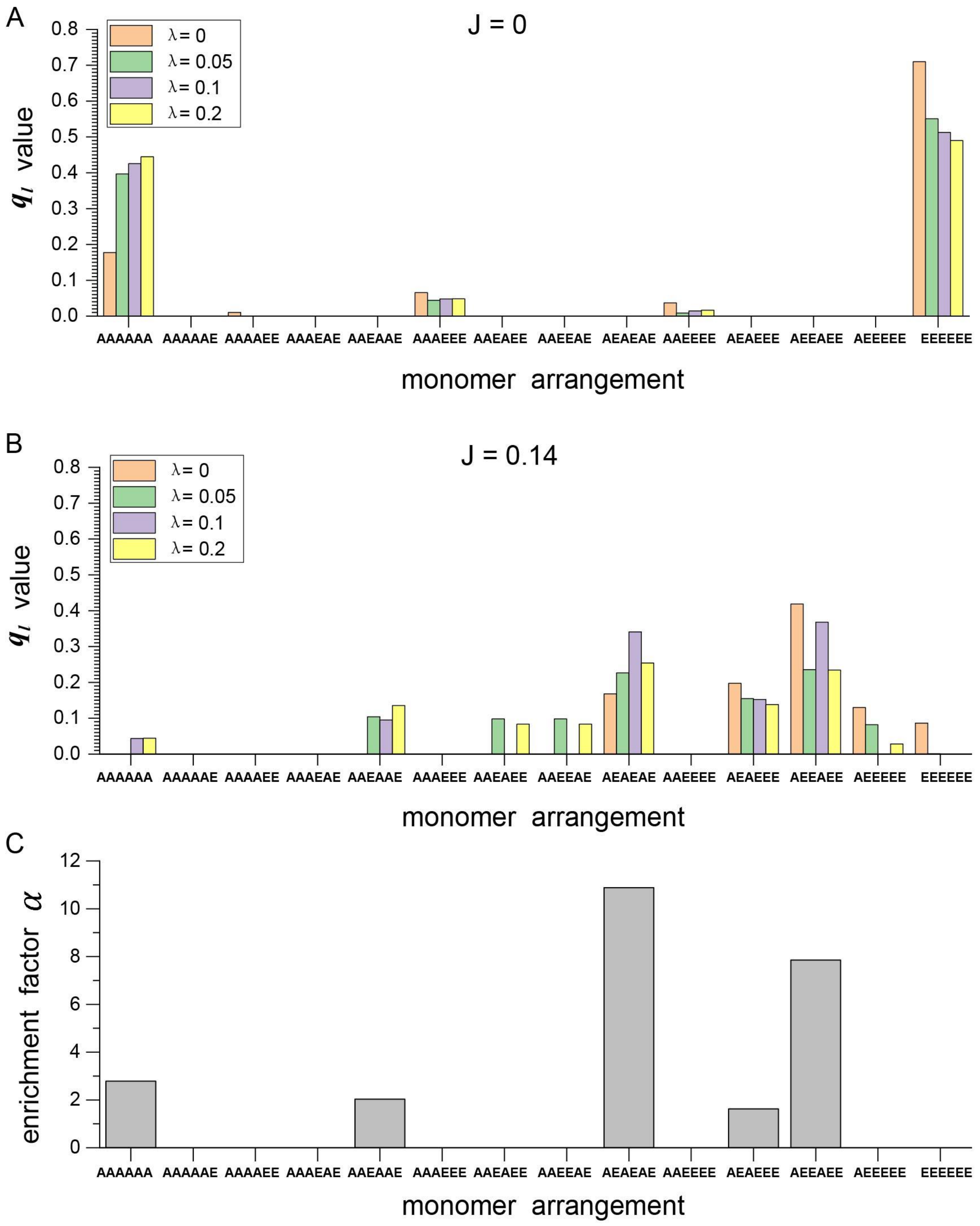


**Supplementary Fig. 5. The monomer arrangement distribution with different choice of $\lambda$ in the case that coupling constant (A) $\mathbf{J}=\mathbf{0}$ or (B) $\mathbf{J}=\mathbf{0.14}$.** The distribution of $q_l$ is insensitive to λ. In the absence of coupling ($\mathrm{J}=0$), the fitting result tells that only pure hexamers exist, which is inconsistent with previous results and definitely not the real case, so the presence of coupling is necessary. (C) The enrichment factor α (the ratio of optimized $q_l$ values when $\lambda=0.1$ in (B) to that under fully-mixed scenario (Supplementary Table 1)) of 14 monomer arrangements. It shows that α >> 1 with the monomer arrangement 'AEAEAE', which indicate that there is a higher probability for two neighboring subunits to have different phosphorylation levels.
